# Alternative reproductive strategies explain asymmetric reinforcement of reproductive isolation in two *Ischnura* damselfly species

**DOI:** 10.1101/2025.05.04.652146

**Authors:** Jesús Ernesto Ordaz-Morales, Alba Leticia Juárez-Jiménez, Miguel Stand-Pérez, Luis Rodrigo Arce-Valdés, Andrea Viviana Ballen-Guapacha, Jesús Ramsés Chávez-Ríos, Adolfo Cordero-Rivera, Rosa Ana Sánchez-Guillén

## Abstract

Theoretical and empirical studies on reinforcement have been a central focus in speciation research. Despite its significance, reinforcement in polymorphic species has received little attention, even though morphs often exhibit differences in behavior and reproductive capacity, which could drive asymmetric reinforcement. In this study, we tested for asymmetric reinforcement between morphs in the polymorphic damselflies *Ischnura elegans* and *Ischnura graellsii*, in which female morphs exhibit alternative reproductive strategies and males show a preference for the gynochrome morph. These species have formed two independent hybrid zones where reinforcement has strengthened a mechanical barrier and reproductive character displacement has shaped mating-related structures. We compared the strength of five reproductive barriers between female morphs to assess the interplay between female-limited color polymorphism and reinforcement. We detected asymmetric reinforcement in premating and postmating barriers between morphs in both hybrid zones, likely driven by their different reproductive strategies. Additionally, we observed weakening of oviposition and fertility barriers probably associated with gene flow and purging of incompatibilities. Our findings highlight that inherent asymmetry in reproductive isolation, caused by differences in female morph reproductive strategies, drive asymmetric reinforcement between morphs. Future studies should explore the cascading effects of this process and its implications for morph speciation.

## INTRODUCTION

Since the 19^th^ century, evolutionary biologists have been interested in understanding the mechanisms driving the emergence of Reproductive Isolation (RI) between species. During speciation, complete RI occurs when reproductive barriers accumulate to prevent genetic exchange (Sobel & Chen, 2014; Westram et al., 2022). However, these barriers evolve at different rates, leading to variation in their relative importance across taxa (Coyne & Orr, 2004). In some taxa, prezygotic barriers—such as habitat or mechanical isolation—are the primary drivers of RI (e.g., Schwarz & McPheron, 2007; Matsubayashi & Katakura, 2009; Sánchez-Guillén et al., 2012; Arce-Valdés et al., 2024). In others, postzygotic barriers—such as hybrid inviability or sterility caused by Bateson-Dobzhansky-Muller (BDM) incompatibility— play a dominant role by reducing the fitness of hybrids (Suni & Hopkins, 2018; Christie & Strauss, 2019; Jiménez-López et al., 2023).

Reinforcement is a widespread speciation process in which natural selection strengthens prezygotic isolation to prevent the production of unfit hybrids, thereby reducing gene flow between diverging heterospecific populations (Hopkins, 2013; Baiz et al., 2019; Coyne & Orr, 2004; St. John & Fuller, 2021; Yukilevich, 2021). This process can also lead to cascade reinforcement, where selection for increased RI in heterospecific interactions indirectly enhances isolation between sympatric and allopatric conspecific populations (Yukilevich & Aoki, 2016; Anderson et al., 2023). Reinforcement often leads to asymmetries in prezygotic isolation between reciprocal heterospecific crosses (Liou & Price, 1994; Hollander et al., 2018) because it typically strengthens prezygotic barriers more rapidly in the cross direction that produces the least fit hybrids (Yukilevich, 2012). One possible explanation for this pattern is that hybrids from reciprocal crosses often differ in viability or fertility— a phenomenon known as “Darwin’s corollary”. This asymmetry arises due to unidirectional accumulation of BDM incompatibilities, often involving sex-chromosomes or maternal effects (Turelli & Moyle, 2007). However, the purging of these incompatibilities in sympatry driven by gene flow between the species, can lead to a weakening of postzygotic barriers (Turelli et al., 2014; Coughlan & Matute, 2020; Xiong & Mallet, 2022) resulting in a reduction of reproductive isolation over time. While reinforcement typically strengthens prezygotic isolation, these dynamics illustrate how gene flow and the purging of BDM incompatibilities can, in some cases, lead to a weakening of postzygotic barriers, complicating the patterns of reproductive isolation between closely related species.

Despite significant progress in understanding the asymmetric patterns emerging from reinforcement, little is known about how this process operates in species with alternative reproductive strategies, where individuals within the same species exhibit distinct mating tactics or morphs. In these cases, RI may vary among morphs due to mechanisms such as ecological segregation (Anthony et al., 2008; Whitney et al., 2018a,b), assortative mating driven by male or female mate choice (Egger et al., 2010) and sex recognition (Powell & Uy, 2023). These differences can lead to asymmetric RI though some exceptions exist (see Pérez-Barros et al., 2011). Most studies on reinforcement in polymorphic species assume that alternative phenotypes function as barriers to heterospecific matings, either due to alternative reproductive strategies or assortative mating favoring conspecifics (Kirkpatrick & Servedio, 1999; Coyne & Orr, 2004; Yukilevich & True, 2006; Nosil & Yukilevich, 2008). However, the extent to which reinforcement actively drives these patterns remains unclear. While some studies have demonstrated reinforcement-driven assortative mating—such as in fishes (Pierotti & Seehausen, 2007) and butterflies (Kronforst et al., 2007)—others suggest that alternative processes, such as reproductive interference, may also contribute, as observed in snails (Hollander et al., 2018). Additionally, cases where reinforcement has been proposed as a potential driver of reproductive character displacement, such as in frogs (Richards-Zawacki & Cummings, 2011), often lack direct tests of reinforcement itself. Understanding how reinforcement interacts with reproductive polymorphisms is therefore essential to disentangle the selective forces shaping RI in these systems.

In this study, we investigate the interplay between polymorphism and reinforcement in two polymorphic damselfly species, *Ischnura elegans* and *Ischnura graellsii,* which are currently undergoing asymmetric reinforcement (Arce-Valdés et al., 2024; Ballén-Guapacha et al., 2024). Both species are non-territorial damselflies that rely on mechanical-tactile cues for species and mate recognition (Robertson & Paterson, 1982), and are ecologically, morphologically, and genetically similar (Monetti et al., 2002; Sánchez-Guillén et al., 2005). *Ischnura elegans* and *I. graellsii* females are polymorphic, exhibiting three mature color morphs: one androchrome morph and two gynochrome morphs (infuscans and aurantiaca). Androchrome and gynochrome females of *I. elegans* and *I. graellsii* exhibit alternative reproductive strategies. For instance, androchrome females of *I. elegans* mimic males in color, morphology, and behavior (Sánchez-Guillén et al., 2005; Van Gossum et al., 2005; Cordero-Rivera & Sánchez-Guillén, 2007; Sánchez-Guillén et al., 2017). In *I. elegans*, males prefer gynochrome females, and gynochrome females exhibit alternative reproductive strategies by mating more frequently and investing more in fecundity compared to androchrome females (Cordero-Rivera & Sánchez-Guillén, 2007; Sánchez-Guillén et al., 2017). Similarly, in *I. graellsii,* males show a preference for gynochrome females (Cordero, 1989)but gynochrome females show lower fecundity compared to the androchrome females, particularly when the operational sex ratio is highly biased toward males (Galicia-Mendoza et al., 2017). These polymorphic traits make these species an excellent model to explore the interaction between alternative reproductive strategies and reinforcement.

These species have established two hybrid zones in Spain—the Northwest (NW) hybrid zone and the Northcentral (NC) hybrid zone—due to the expansion of *I. elegans* into North central and Western Spain (Sánchez-Guillén et al., 2005, 2023). In these hybrid zones, several patterns have been observed: i) high frequency of androchrome morphs in both species (Sánchez-Guillén et al., 2005); ii) variable introgression and hybridization at local and regional scales: bidirectional introgression and hybrid swarm in the Northwest hybrid zone and unidirectional introgression towards *I. graellsii* in the Northcentral hybrid zone (Sánchez-Guillén et al., 2023); iii) asymmetric RI between reciprocal crosses (Sánchez-Guillén et al., 2012; Arce-Valdés et al., 2024); and iv) reproductive character displacement (RCD) in male caudal appendages of both species and female prothorax of *I. elegans* (Ballén-Guapacha et al., 2024) by reinforcement of a mechanical premating barrier in the heterospecific crosses between *I. graellsii* males and *I. elegans* females (Arce-Valdés et al., 2024). In *I. elegans,* the detection of RCD in the male caudal appendages and female prothorax as consequence of reinforcement has been taken as an evidence of the coevolution of these reproductive structures due to lock-and-key mechanism (c.f. Dufour, 1844) driven by female choice of conspecific males (Ballén-Guapacha et al., 2024).

The main aim of our study was to investigate the interplay between polymorphism and reinforcement in two female color morphs of *I. elegans* and *I. graellsii* across the two hybrid zones. Specifically, we compare the strength of five prezygotic barriers between androchrome and gynochrome morphs in both species from allopatry, and both hybrid zones. We predict that in allopatry androchrome females will exhibit higher levels of RI in heterospecific crosses due to their morphological similarity to males in prothorax shape, i.e., the structure involved in the first contact point between male and females during copulation (Ballén-Guapacha et al., 2024), their reluctance to accept matings, and male mating preference for gynochrome females (Cordero, 1989; Cordero-Rivera & Sánchez-Guillén, 2007; Galicia-Mendoza et al., 2017; Sánchez-Guillén et al., 2017), while in sympatry both female morphs will exhibit similar levels of RI due to reinforcement of the gynochrome females. In addition, we examined whether male-mimicry in androchrome females extends beyond their morphological and behavioral traits by comparing the success male-male tandem attempts with both male-androchrome and male-gynochrome tandem attempts in both *I. elegans* and *I. graellsii*. This hybrid system provides a unique opportunity to empirically investigate the early stages of speciation, offering valuable insights into evolutionary processes in hybridizing species.

## MATERIALS AND METHODS

### Hybrid zones description

The Spanish sympatric zone consists of at least two independent hybrid zones: the Galician hybrid zone, hereafter referred to as the Northwest (NW) hybrid zone, and the Northcentral and Mediterranean hybrid zone (Fig. 1). We focused our study on the NW hybrid zone, and on a small zone in the Northcentral and Mediterranean hybrid zone, hereafter referred to as the NC hybrid zone. In the Spanish sympatric zone, the proportions of species and hybrids vary among populations and zones, as well as the frequency of female morphs, which contrast with those observed in the allopatric distribution of both species (Sánchez-Guillén et al., 2005, 2023). For instance, researchers have reported a relatively stable frequency of androchrome females in *I. elegans* across its allopatric distribution, with an increasing trend toward northern latitudes (Van Gossum et al., 2005; Hammers & Van Gossum, 2008; Cordero-Rivera et al., 2024), with some populations exhibiting exceptional cases where the relative frequency reaches approximately 76% (Hammers & Van Gossum, 2008). In sympatry, however, the relative frequency of androchrome females varies widely among populations, ranging from 8% to 90% (Cordero-Rivera & Sánchez-Guillén, 2007; Ordaz-Morales, 2022). The NW hybrid zone is characterized by greater bidirectional introgression, while the NC hybrid zone is characterized by lower unidirectional introgression towards *I. graellsii* (Sánchez-Guillén et al., 2023). Furthermore, the degree of genetic differentiation between species in the NW hybrid zone is lower than that found in the NC hybrid zone, suggesting that the former is less exposed to gene flow from the allopatric distribution of *I. elegans* (Sánchez-Guillén et al., 2023).

**Figure 1.**
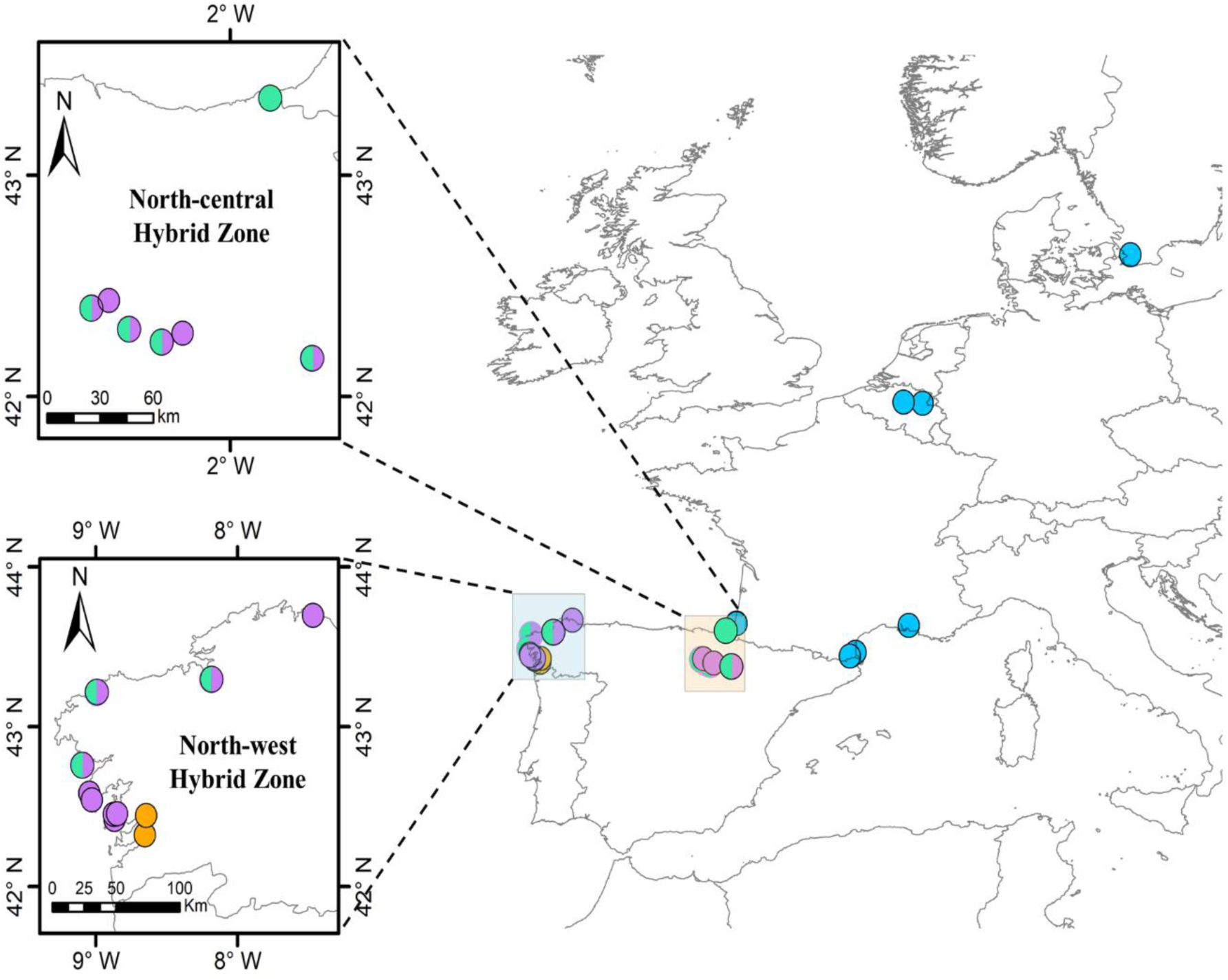
Sampling localities in allopatry and hybrid zones of *I. elegans* and *I. graellsii*. Green square represents zoomed-in areas in the NW hybrid zone. Beige square represents zoomed-in areas in the NC hybrid zone. Blue circles indicate sampled localities for allopatric *I. elegans*, while orange circles show sampled localities for allopatric *I. graellsii*. Purple circles denote sympatric localities where just *I. graellsii* was sampled, and green circles denote sympatric localities where just *I. elegans* was sampled. Bicolor circles indicate sympatric localities where both species were sampled.

### Field sampling

In 2023 and 2024 we sampled two *I. graellsii* populations (Louro and Xuño), two *I. elegans* populations (Doniños and Laxe), and one population with both species (Brañas) from the NW hybrid zone. We also sampled two allopatric *I. graellsii* populations: Riomaior and Cachadas. Similar data from allopatric populations, NC hybrid zone and NW hybrid zone are available from three previous studies (Sánchez-Guillén et al., 2012; Sánchez-Guillén et al., 2023; Arce-Valdés et al., 2024). See Table S1 for details on localities and number of mating pairs from this study as well as from previous studies. To summarize, in this study we produced 487 mating pairs, which were added to the previous 406 mating pairs (n=893). In total, we analysed 119 conspecific mating pairs of both species from 10 localities from allopatry, 39 conspecific mating pairs of both species from four localities in the NC hybrid zone, and 119 conspecific mating pairs of both species from nine localities in the NW hybrid zone, 81 heterospecific mating pairs from 11 localities in allopatry, 60 heterospecific mating pairs from five localities in the NC hybrid zone and 470 heterospecific mating pairs from 11 localities in NW hybrid zone. Details on conspecific male-female crosses are provided in Table S2. Data on mechanical isolation was also collected in male-male interactions. To this end, we sampled six populations from the NW hybrid zone, including 51 male-male interactions performed by 39 *I. elegans* males, and 138 male-male interactions performed by 82 *I. graellsii* males.

### Rearing in the laboratory and mating trials

We implemented a modified rearing protocol (see Van Gossum et al., 2003; Sánchez-Guillén et al., 2012). We maintained last-instar larvae and tenerals in the laboratory until they reached sexual maturity. Adults were marked with a unique code on the wing for identification and males, virgin females and mated females were kept separated in in 50 x 50 x 50 cm wooden insectaries (Van Gossum et al., 2003). For mating trials, we followed the method outlined by Sánchez-Guillén et al., (2012). We conducted several conspecific and heterospecific mating trials to achieve all possible crosses between *I. elegans* and *I. graellsii*. This involved placing between one to fourteen males and one to ten females in a wooden insectary during their reproductive activity period. The numbers of males and females, as well as the proportion of females morph per insectary were determined by the availability of sexually mature individuals per day. Given that males of both species differ in the time of maximum reproductive activity (from 09:00h to 14:00h for *I. elegans* and from 12:00h to 17:00h for *I. graellsii*), each mating trial started based on the male species and was performed for a maximum five hours or until the possible mating was achieved. Once the mating finished, we isolated the mated females and provided them with the conditions to oviposit (Van Gossum et al., 2003; Sánchez-Guillén et al., 2012). Mated females were allowed to oviposit each day from 15:00h to 08:00h of the following day until they died. Larvae were reared up to adulthood following standardized protocols (Van Gossum et al., 2003; Sánchez-Guillén et al., 2012), and mating trials were repeated in the F_2_ generation. Finally, since tandem formation is possible between males, we also measured the mechanical isolation in male-male interactions and compared it to male-female interactions with each morph.

### Reproduction in *Ischnura* and measurement of reproductive isolation

In damselflies, mating begins when a male successfully grasps the female by her prothorax using his caudal appendages, achieving the ’tandem position’ (Robertson & Paterson, 1982). The females may accept copulation bending her abdomen and the genitalia engage, forming the so called ’wheel position’ (Cordero, 1989). Nevertheless, given that damselflies rely on mechanosensory cues for mate recognition, it is possible to observe the tandem position in male-male interactions (e.g. Beatty et al., 2015; Khan & Herberstein, 2021). After copulation, the female lays eggs until her egg reserve is depleted.

We used each male-female pair or mated female as the unit of observation to evaluate five reproductive barriers (Table 1), classified into two premating barriers and three postmating barriers, for conspecific and heterospecific crosses (prezygotic isolation). These barriers were: i) prevention of tandem formation (premating mechanical barrier), ii) prevention of genitalia engagement and the formation of the wheel position (premating mechanical-tactile barrier), iii) prevention or reduction of female oviposition (postmating oviposition barrier), iv) reduction of female fecundity (postmating fecundity barrier), and v) reduction of female fertility (postmating fertility barrier).

**Table 1.** Summary of the formulas used to estimate absolute reproductive isolation (RI) and their descriptions as included in the statistical analysis (GLMs and post-hoc) for each reproductive barrier. RI was estimated by adapting the general formula of Coyne and Orr (1989) as follows : 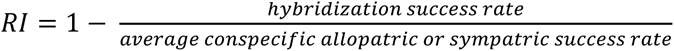

| Reproductive barrier | GLM response variable description | Absolute reproductive isolation formula |
| --- | --- | --- |
| **Premating: estimated using unique interacting reproductive male-female pairs** |  |  |
| I. Mechanical | Binomial: total of successful tandems vs total of tandem attempt | $RI = 1 - \frac{\frac{\text{Total succesful tandems}}{\text{Total tandem attempts}}}{\text{Conspecific control}}$ |
| II. Mechanical - Tactile | Binomial: total of successful matings vs total of tandems | $RI = 1 - \frac{\frac{\text{Total succesful matings}}{\text{Total mating attempts}}}{\text{Conspecific control}}$ |
| **Postmating: estimated using isolated mated females, i.e., which successfully formed copulation positions with a single male (n)** |  |  |
| I. Oviposition | Binomial: number of mated females that laid eggs vs<br>number of mated females that did not lay eggs | $RI = 1 - \frac{\frac{\text{Mated females that laid eggs}}{\text{number of mated females}}}{\text{conspecific control}}$ |
| II. Gametic (Fecundity) | Gaussian: average number of eggs laid during the first<br>three clutches per female | $RI = 1 - \frac{(\sum_{i=1}^n \text{eggs per clutch index})/n}{\text{Conspecific control}}$ |
| III. Gametic (Fertility) | Binomial: number of fertile eggs vs number of infertile eggs | $RI = 1 - \frac{\left(\frac{\sum_{i=1}^n \text{fertile eggs}}{\text{total laid eggs}}\right)/n}{\text{Conspecific control}}$ |

**Table 2.** Absolute reproductive isolation, sample size (in parentheses), and 95% confidence interval (CI) for each reproductive barrier measured in heterospecific crosses in allopatry, NW and NC hybrid zones. For mechanical and mechanical-tactile barriers, the sample size corresponds to the number of tandem attempts and successful tandems, respectively, for each cross direction. E × G represents *I. elegans* x *I. graellsii*, G × E represents *I. graellsii* x *I. elegans*. A denotes androchrome females, and G denotes gynochrome females.

| Cross<br>(♂×♀) | Morph | Mechanical |  | Mechanical-Tactile |  | Oviposition |  | Fecundity |  | Fertility |  |
| --- | --- | --- | --- | --- | --- | --- | --- | --- | --- | --- | --- |
|  |  | mean | CI | mean | CI | mean | CI | mean | CI | mean | CI |
| Allopatry |  |  |  |  |  |  |  |  |  |  |  |
| E × G | A | 0.198 (15) | -0.634 to 0.627 | -0.322 (8) | -1.692 to 0.444 | -0.104 (3) | -0.270 to 0.000 | 0.906 (3) | 0.778 to 0.965 | 1.000 (3) | 1.000 to 1.000 |
|  | G | -0.418 (59) | -1.133 to 0.126 | 0.121 (48) | -0.329 to 0.453 | 0.293 (28) | = 0.095 to 0.495 | 0.717 (18) | 0.101 to 0.972 | 0.934 (18) | 0.585 to 1.000 |
| G × E | A | 0.624 (4) | -0.270 to 1.000 | -0.511 (1) | -3.337 to 1.000 | -0.104 (1) | -0.268 to 0.000 | -0.543 (1) | -1.322 to -0.104 | -0.148 (1) | -0.486 to 0.047 |
|  | G | -0.743 (12) | -1.678 to -0.063 | 0.168 (12) | -0.266 to 0.521 | -0.100 (6) | -0.174 to -0.044 | -0.481 (6) | -2.475 to 0.195 | 0.066 (6) | -0.306 to 0.692 |
| Northcentral hybrid zone |  |  |  |  |  |  |  |  |  |  |  |
| E × G | A | 0.313 (32) | -1.155 to 0.763 | 0.192 (13) | -1.424 to 1.000 | -0.059 (3) | -0.250 to 0.000 | -2.057 (3) | -9.398 to 0.358 | -0.305 (3) | -2.512 to 0.833 |
|  | G | 0.213 (32) | -0.658 to 0.660 | -0.077 (18) | -1.294 to 0.542 | 0.027 (14) | -0.143 to 0.214 | 0.681 (13) | 0.081 to 0.968 | 0.670 (13) | -0.102 to 1.000 |
| G × E | A | 0.324 (5) | -1.800 to 1.000 | -0.312 (2) | -5.000 to 1.000 | -0.059 (2) | -0.250 to 0.000 | -1.487 (2) | -10.558 to -0.123 | 0.897 (2) | 0.707 to 1.000 |
|  | G | 0.696 (23) | 0.280 to 0.926 | -0.108 (5) | -3.374 to 0.818 | -0.048 (4) | -0.173 to 0.000 | 0.505 (4) | -0.533 to 0.886 | -0.215 (4) | -1.527 to 0.982 |
| Northwest hybrid zone |  |  |  |  |  |  |  |  |  |  |  |
| E × G | A | 0.455 (164) | -0.392 to 0.769 | 0.132 (44) | -2.117 to 1.000 | 0.039 (18) | -0.054 to 0.172 | 0.401 (17) | -0.503 to 0.833 | 0.106 (17) | -0.213 to 0.397 |
|  | G | 0.286 (382) | 0.007 to 0.506 | 0.012 (161) | -0.435 to 0.338 | 0.032 (100) | -0.041 to 0.100 | 0.403 (93) | 0.090 to 0.631 | 0.250 (93) | 0.107 to 0.391 |
| G × E | A | 0.966 (177) | 0.885 to 1.000 | 0.394 (3) | -4.100 to 1.000 | -0.018 (2) | -0.074 to 0.000 | 0.217 (2) | -0.222 to 0.494 | 0.220 (2) | 0.055 to 0.265 |
|  | G | 0.947 (351) | 0.904 to 0.977 | 0.189 (11) | -1.167 to 0.777 | -0.041 (7) | -0.099 to 0.000 | 0.504 (7) | -0.476 to 0.906 | 0.262 (7) | -0.131 to 0.881 |

The mechanical barrier was assessed by counting the number of tandem attempts that each pair performed before the male successfully grasped the female by her prothorax. Furthermore, the mechanical barrier was assessed in a similar manner in conspecific male-male interactions. The mechanical-tactile barrier compared pairs in which the female bent her abdomen and genitalia engaged with those in which the female bent her abdomen, but the mating organs were mechanically incompatible. The oviposition barrier was measured by comparing the number of mated females that laid eggs to those that did not. Thus, the mechanical, mechanical-tactile, and oviposition barriers were treated as binomial variables. The fecundity barrier was measured as the average number of eggs laid per mated female and modeled using the Gaussian distribution (eggs per clutch index). Finally, the fertility barrier was measured as a binomial variable by comparing the number of fertile eggs to infertile eggs.

### Estimation of absolute and relative strength of the reproductive barriers

Our primary objective was to compare the amount of gene flow in heterospecific crosses between female color morphs of both species. To this end, to estimate the absolute strength of each reproductive barrier we applied the formulae described by previous studies in this hybrid system (Sánchez-Guillén et al., 2023; Arce-Valdés et al., 2024) which are a modified version of the Coyne and Orr (1989) formula to estimate the absolute strength of each reproductive barrier:

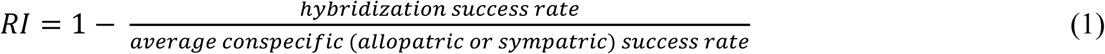

Conspecific crosses of both species and female morphs (androchrome and gynochrome) were used as controls to estimate the strength of the reproductive barriers in heterospecific crosses (Sánchez-Guillén et al., 2012; Barnard et al., 2017; St. John & Fuller, 2021, Table S2). Specifically, we assigned a specific control to each morph, species, and zone (allopatric, NC or NW hybrid zone) meaning that the control consisted of the conspecific cross for that particular morph, species and zone. These controls allow us to compare between heterospecific and conspecific crosses, while also accounting for potential differences in conditions between the species natural environments and experimental setting (Sobel & Chen, 2014). By applying the conspecific control values to our formula, the absolute strength of each reproductive barrier can range from −1 to 1. Values above 0 indicate stronger reproductive isolation in heterospecific crosses compared to conspecific crosses for a given female morph. Conversely, values below 0 indicate that the barrier facilitates hybridization. We also estimated the cumulative contribution of each reproductive barrier to total reproductive isolation by applying the multiplicative function of individual components developed by Coyne and Orr (1989; 1997) and later modified by Ramsey et al. (2003) to include any number of reproductive barriers (Sobel & Chen, 2014). The cumulative contribution (*CC*) of a component (reproductive barrier) to the RI at a stage *n* was estimated with the following formula:

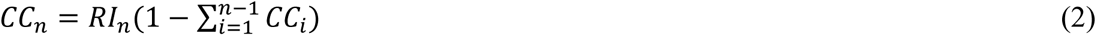

In this equation, *RI_n_* represents the absolute RI of each barrier at stage *n.* At *CC_1_* we used the absolute RI of the mechanical barrier which is the earliest barrier measured.

#### Confidence interval around RI estimates

We assessed uncertainty around the mean estimates of RI using a bootstrapping approach, following the method of Christie & Strauss (2019). We resampled the raw data with replacement and calculated RI 10,000 times, as described above, then computed 95% confidence intervals in R 4.3.1 (R Core Team 2023). When the 95% confidence interval included zero, we inferred that RI was zero, indicating that the measured barrier performed similarly to conspecific controls. Additionally, we used the overlap in the estimated upper and lower CI to determinate the asymmetry in the strength of each reproductive barrier between female morphs.

### Reinforcement or weakening of the reproductive barriers

To investigate asymmetries in the reinforcement or weakening of RI between female morphs we used Generalized Lineal Models (GLMs) to assess the association between each reproductive barrier and the cross type (*I. elegans* males and *I. graellsii* females vs *I. graellsii* males and *I. elegans* females), distribution (allopatry vs NW hybrid zone vs NC hybrid zone), female morph (androchrome vs gynochrome) and an interaction variable (crosses*distribution*morph) as predictor variables. We built a unique global model including each predictor variable using the *glm* function in R 4.3.1 (R Core Team 2023). Thus, our global model for each barrier was formulated as follows: *Isolation ∼ Cross + Distribution + Morph + Interaction.* We fitted the mechanical, mechanical-tactile, oviposition and fertility barriers using the binomial distribution. Additionally, we fitted the fecundity barrier (eggs per clutch index) using the Gaussian distribution. To ensure model adequacy and goodness-of-fit of each constructed model we simulated its residuals using the function *simulateResiduals* of the *DHARMa* 0.4.6 library (Hartig, 2024).

Our *post hoc* analysis to assess differences between hybrid zones *versus* allopatry for each female morph consisted of a *Wald z*-test for the binomial variables and the *Tukey* test for the Gaussian variables. These *post hoc* tests were conducted by estimating the marginal means and contrasting the interaction variable of the global model using the *emmeans* and *contrast* functions from *emmeans* 1.10.4 library (Lenth, 2025). Finally, we employed the sequential Holm-Bonferroni correction (Bonferroni, 1936; Holm, 1979) to reduce risk of Type I error due to multiple comparisons. Note that, when we observed that a given barrier behaved statistically similarly across zones (i.e., no significant differences were detected between hybrid zones) and the statistical values in the zone-specific analyses were close to significance before the sequential Holm-Bonferroni correction, we combined the data from both hybrid zones into a sympatric category. This allowed us to compare the RI of both morphs in an allopatry vs. sympatry framework to enhance the statistical power of our analysis. Finally, we implemented a two-proportion *z*-test to compare male-male mechanical isolation against male-female mechanical isolation for each female morph.

## RESULTS

For clarity, the main document presents the analyses by zones (allopatry, NC, and NW hybrid zones). Additional analyses that combine data from both hybrid zones are included in the supplementary material (Tables S4–S5, Figs. S3–S5).

### Reproductive isolation by female morphs across hybrid zones

#### Androchrome female morphs in heterospecific crosses

In allopatry, *I. elegans* females exhibited weak reproductive isolation (RI = −0.012; Fig. 2), with mechanical isolation (RI = 0.624; Fig. S1A) being the main significant premating barrier. However, postmating barriers, such as oviposition (RI = −0.104; Fig. S1C) and fecundity (RI = −0.543; Fig. S1D), significantly reduced this effect. In contrast, *I. graellsii* females displayed complete reproductive isolation (RI = 1.000; Fig. 2), with key significant postmating barriers, such as fecundity (RI = 0.906; Fig. S1D) and fertility (RI = 1.000; Fig. S1E), completely preventing hybrid larval production.

**Figure 2.**
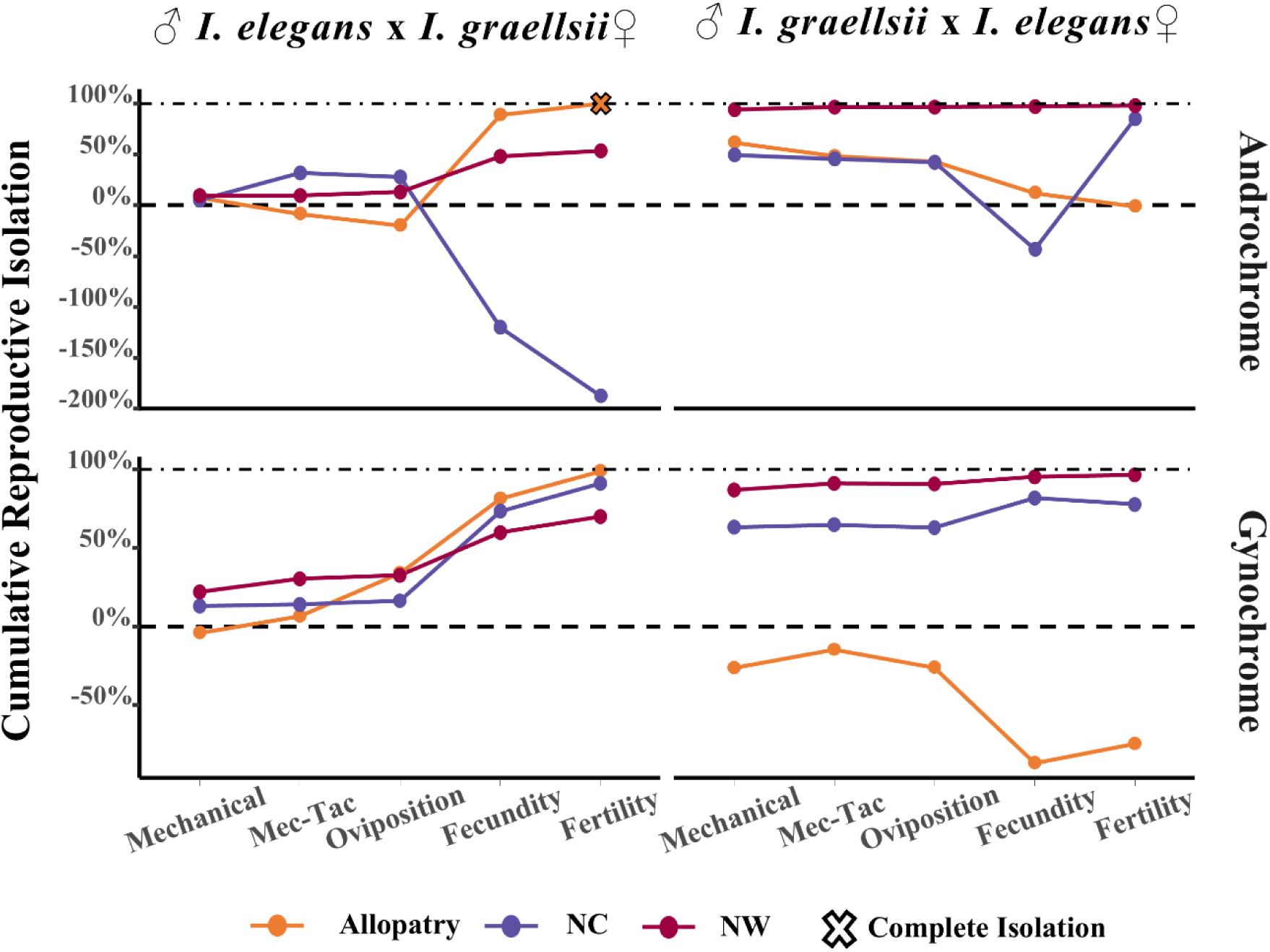
Cumulative isolation in both reciprocal crosses between *I. elegans* and *I. graellsii*. Upper panels display cumulative reproductive isolation for androchrome females, while lower panels show cumulative reproductive isolation for gynochrome females in allopatry (orange dots and line), NW hybrid zone (red dots and line), and NC hybrid zone (purple dots and line). A cross indicates complete reproductive isolation and a lack of gene flow between species in that cross direction. Horizontal dashed line indicates the threshold between reproductive isolation (above the line) and hybridization (below the line).

In the NC hybrid zone, *I. elegans* females showed an overall high isolation (RI=0.853; Fig. 2), explained by both premating mechanical (RI=0.324 Fig. S1A) and postmating fertility (RI=0.897; Fig. S1E) barriers. Meanwhile, *I. graellsii* females exhibited highly weakened reproductive isolation (RI =-1.874; Fig. 2). The cumulative effect of premating barriers— specifically mechanical (RI=0.313; Fig. S1A) and mechanical-tactile (RI=0.192; Fig S1B)— were the primary barriers limiting hybridization. However, this effect vanished due to weakening postmating barriers such as oviposition (RI=-0.059; Fig. S1C) and fecundity: RI=-2.057; Fig. S1D).

In the NW hybrid zone, *I. elegans* females showed nearly complete isolation (RI= 0.978; Fig. 2). Overall, premating barriers played a dominant role in preventing hybridization (mechanical: RI=0.966; Fig S1A), while postmating barriers contributed to a lesser extent (fecundity RI = 0.217, fertility RI = 0.220; Fig. S1D-E). On the other hand, *I. graellsii* females displayed moderate isolation (RI= 0.698; Fig. 2). Overall, postmating barriers were the primary barriers limiting hybridization (oviposition: RI=0.039, fecundity: RI=0.401, and fertility: RI=0.106; Fig. S1C-E).

#### Gynochrome female morphs in heterospecific crosses

In allopatry, *I. elegans* females exhibited weak reproductive isolation (RI=-0.748; Fig. 2), neither premating mechanical barrier (RI=-0.743; Fig. S1A) nor postmating barriers such as oviposition (RI=-0.100; Fig S1C) and fecundity (RI=-0.481; Fig. S1D) effectively prevented hybridization for this female morph. In contrast, *I. graellsii* females displayed nearly complete reproductive isolation (RI=0.988, Fig. 2), with key significant postmating barriers—oviposition (RI=0.293; Fig. S1C), fecundity (RI=0.717 Fig. S1D), and fertility (RI=0.934; Fig. S1E)—impeded hybridization in this cross direction.

In the NC hybrid zone, *I. elegans* females showed overall high reproductive isolation RI =0.778; Fig. 2), with mechanical isolation (RI=0.696; Fig. S1A) as the primary barrier limiting hybridization. Meanwhile, *I. graellsii* females exhibited a stronger reproductive isolation (RI=0.912, Fig. 2), with both premating (mechanical isolation: RI=0.213; Fig. S1A), and postmating barriers—oviposition (RI=0.027; Fig. S1C), fecundity (RI=0.681; Fig. S1D), and fertility (RI=0.670; Fig. S1E)—contributed to reproductive isolation.

In the NW hybrid zone, *I. elegans* females showed high reproductive isolation (RI= 0.966; Fig. 2). Overall, premating barriers played a dominant role in preventing hybridization (mechanical: RI=0.947; Fig. S1A), while postmating barriers contributed to a lesser extent (fecundity RI = 0.504, fertility RI = 0.262; Fig. S1D–E). On the other hand, *I. graellsii* females displayed moderate reproductive isolation (RI= 0.698; Fig. 2). Both premating (mechanical: RI=0.286, and mechanical-tactile: RI=0.012; Fig. S1A-B) and postmating barriers (fecundity: RI=0.403, and fertility: RI=0.250; Fig. S1C-E) contributed similarly to preventing hybridization.

### Reinforcement of the reproductive barriers by female morphs

Statistical details on the *post hoc* contrast tests between allopatry and hybrid zones were summarized in the Supplementary Table S4.

#### Androchrome female morphs in heterospecific crosses

When comparing allopatry and the NC and NW hybrid zones, for the *I. elegans* females, we detected a significant strengthening of the fertility barrier (NC: *p* < 0.001; NW: *p* = 0.023; Fig. 3C;). In contrast, for *I. graellsii* females, we detected, in both hybrid zones, a significant weakening of the fertility barrier (NC: *p* = 0.001; NW: *p* < 0.001; Fig. 3C).

**Figure 3.**
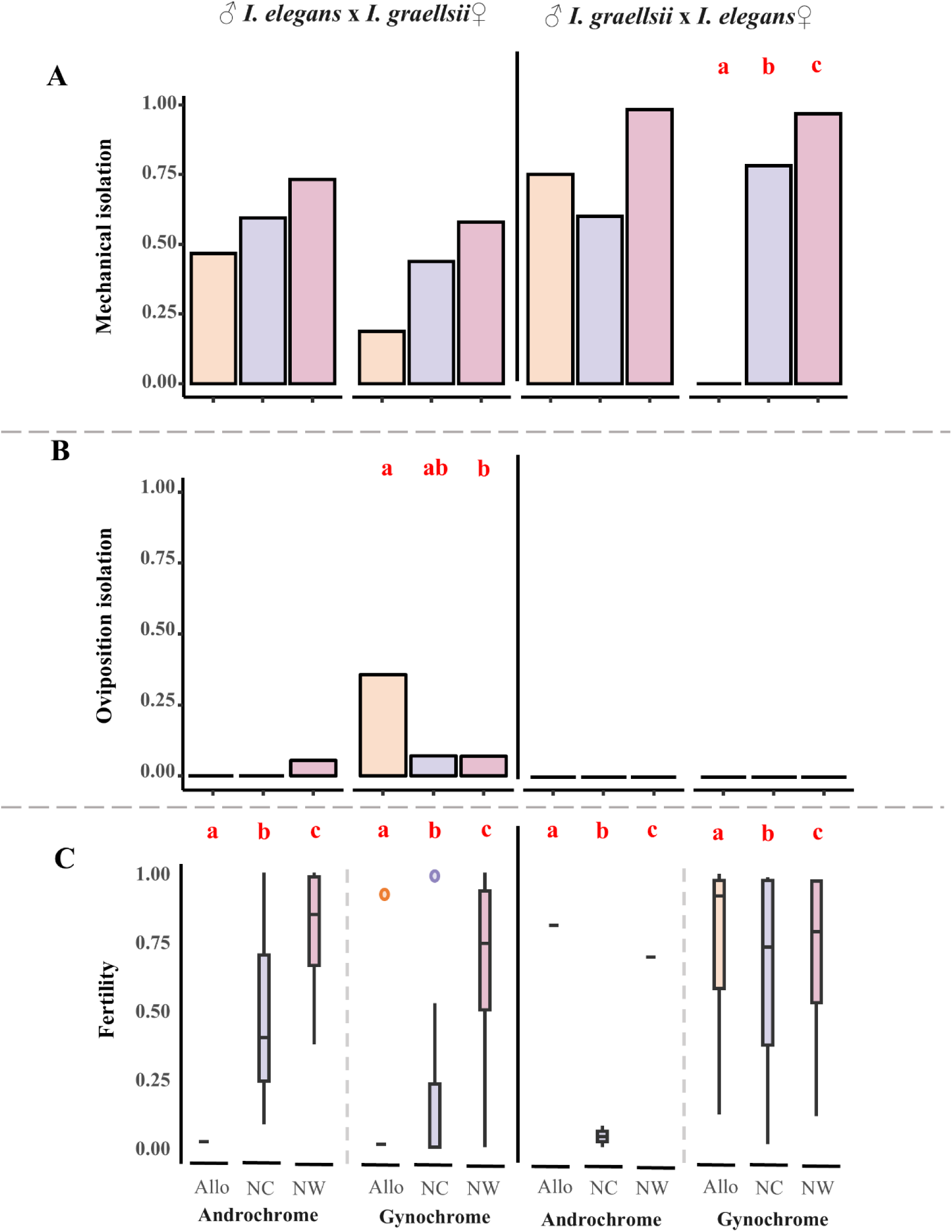
Differences in prezygotic barriers between hybrid zones. Significant differences in three prezygotic barriers for androchrome and gynochrome females in allopatry (beige bar), NC hybrid zone (purple bar), and NW hybrid zone (pink bar). **A)** mechanical isolation, measured as the number of tandems per tandem attempts per heterospecific pair. **B**) oviposition isolation, measured as the number of females that oviposited after mating. **C)** fertility, measured as the ratio of fertile eggs laid by each female. Red bold letters above the barplot, and boxplot indicate statistically significant difference after Holm-Bonferroni correction.

#### Gynochrome female morphs in heterospecific crosses

For the *I. elegans* females when comparing allopatry and the NC and NW hybrid zones, we detected a significant increase in the mechanical (NC: *p* < 0.001; Fig. 3A; NW: *p* < 0.001; Fig 3A) and the fertility barriers (NC: *p* < 0.001; NW: *p* < 0.001; Fig. 3C). Since the fecundity barrier in gynochrome females of *I. elegans* showed similar patterns across zones (Table S4) but marginal statistical significance in the NW hybrid zone (*p* = 0.0085, Holm threshold = 0.0083; Fig. 3A) probably due the small sample size per hybrid zone, we combined both hybrid zones, thus making a comparison between allopatry and sympatry, and thus we detected a significant strengthening of the fecundity barrier (*p* = 0.003, Fig. 4B).

**Figure 4.**
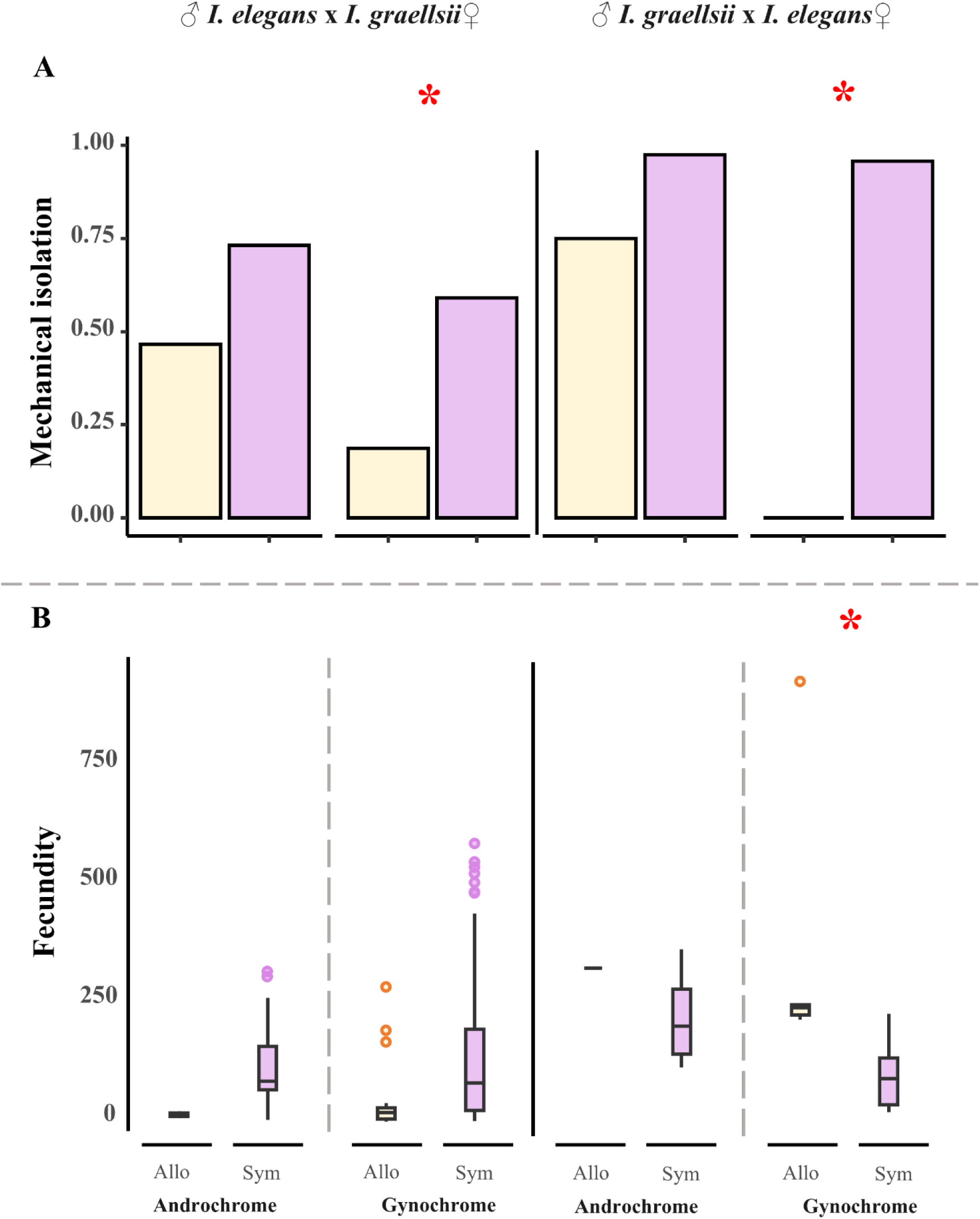
Differences in prezygotic barriers after merging data from both hybrid zones. Significant differences in two prezygotic barriers for androchrome and gynochrome females in allopatry (yellow bar), and sympatry (pink bar). **A)** mechanical isolation, measured as the number of tandems per tandem attempts per heterospecific pair. **B**) fecundity, measured as the number of eggs laid by each female. Red asterisks indicate statistically significant difference after Holm-Bonferroni correction analyzed with the raw data.

In contrast, for *I. graellsii* females, in both hybrid zones, we detected a significant weakening of the fertility barrier (NC: *p* = 0.001; NW: *p* < 0.001; Fig. 3C), and a significant weakening the oviposition barrier, but only in the NW hybrid zone (*p* = 0.003; Fig. 3B). Additionally, gynochrome females of *I. graellsii* females exhibited stronger mechanical barrier when both zones were combined into a single sympatry (*p* = 0.026; Fig. 4A). Analyses by zones showed only marginal significance for the NW zone (*p* = 0.0085, Holm threshold = 0.0083; Fig. 3A).

### Differences in mechanical isolation between males and female morphs

We quantified male–male tandem success and compared it to male-female tandem success in conspecific crosses of each species. In *I. elegans*, males showed low levels of tandem success (0.289) in male-male interactions. As expected, tandem success in male-male interactions was significantly lower than in male-gynochrome interactions (tandem success = 0.595, *X*^2^ = 11.815, df = 1, *p* < 0.001; Fig. 5) but not lower than in male-androchrome interactions (tandem success = 0.484, *X*^2^ = 3.213, df = 1, *p* = 0.073; Fig. 5). Similarly, *I. graellsii* males showed almost null tandem success in male-male interactions (tandem success = 0.043), which was significantly lower than the success observed in interactions with both gynochrome (tandem success = 0.586, *X^2^* = 128.900, df = 1, *p* < 0.001; Fig. 5) and androchrome (tandem success = 0.500, *X*^2^ = 17.242, df = 1, *p* < 0.001; Fig. 5) females.

**Figure 5.**
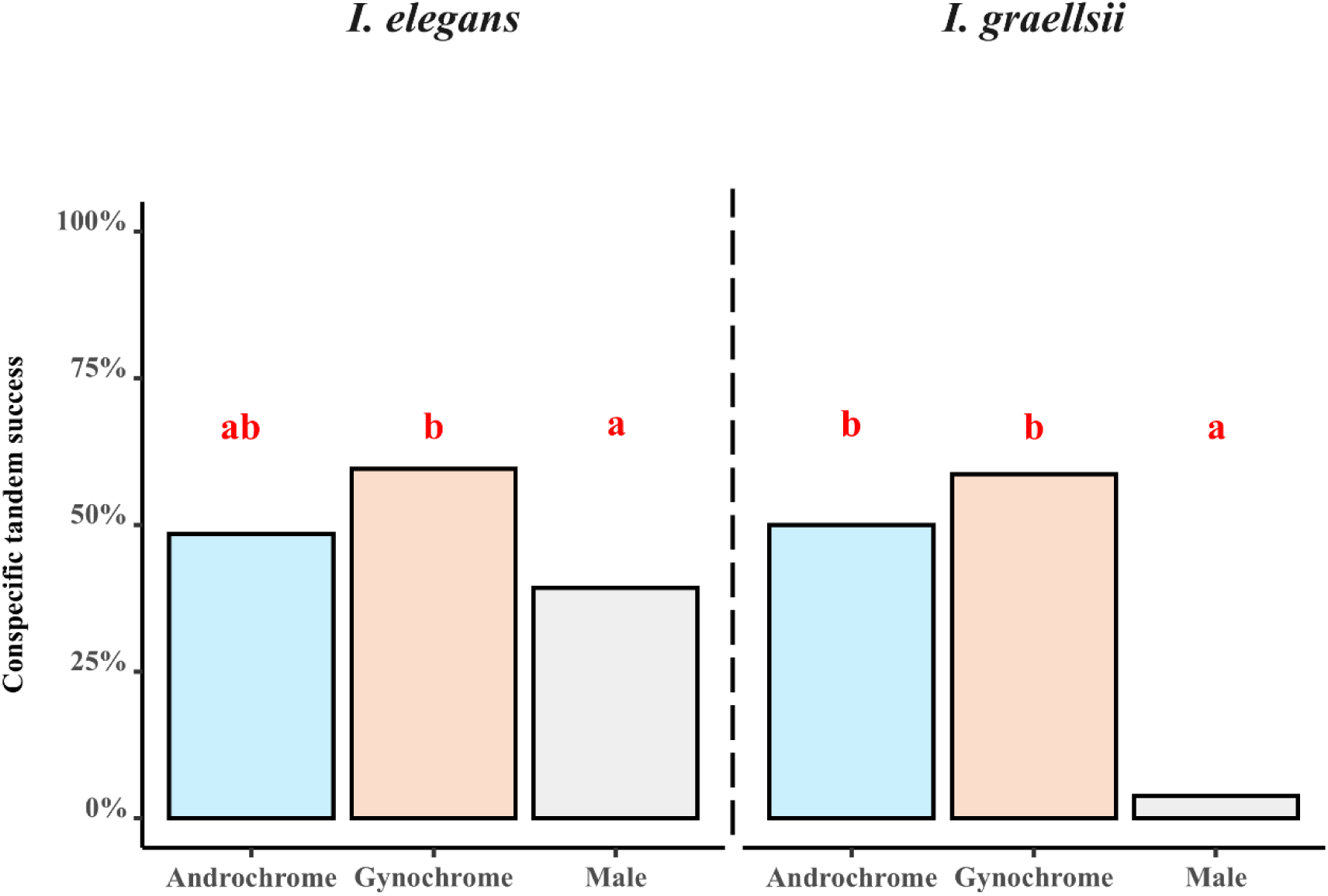
Conspecific tandem success in the NW hybrid zone. Barplot of the tandem success, measured as the number of tandems per tandem attempts in conspecific pairs: male-androchrome (blue bar), male-gynochrome (beige bar) and male-male (grey bar) interactions. Red bold letters above the barplot indicate statistically significant differences.

## DISCUSSION

Our study evaluated the strength of prezygotic barriers in the androchrome and gynochrome morphs of *I. elegans* and *I. graellsii* in two hybrid zones, providing a unique opportunity to examine the interaction between alternative reproductive strategies and reinforcement. Our results indicate that reinforcement of the mechanical barrier acted asymmetrically between female morphs, strengthening only in the gynochrome females of both species. This pattern may be explained by the inherently higher mechanical isolation of the androchrome females due to their reproductive strategy. In contrast, changes in the strength of the gametic barriers followed a symmetric pattern between morphs but differed between species: reinforcement of the fertility barrier occurred in *I. elegans* females, whereas *I. graellsii* females exhibited weakening of the oviposition and fertility barriers.

### Asymmetric reinforcement of the mechanical barrier between female morphs

Our results showed that reinforcement of the mechanical barrier acted asymmetrically between female morphs, strengthening only in the gynochrome females of both species. Although reinforcement in polymorphic species remains poorly studied, available studies support our findings, showing that reinforcement can act asymmetrically between morphs, driven by both intrinsic and extrinsic factors (Liou & Price, 1994; Kirkpatrick & Servedio, 1999; Noor, 1999; Servedio, 2001; Coyne & Orr, 2004; Yukilevich, 2012; Garner et al., 2018; Yukilevich et al., 2024). In fishes (Pierotti & Seehausen, 2007; Egger et al., 2010), walking-sticks (Nosil et al., 2003), and poison frogs (Richards-Zawacki & Cummings, 2011; Yang et al., 2016), assortative mating reinforces isolation between morphs. For instance, in the walking stick *Timema cristinae,* morphs adapted to different host plants exhibit reinforced mate discrimination against non-local males to avoid producing maladaptive hybrids (Sandoval, 1994; Nosil et al., 2003). Extrinsic factors also play a role, as seen in honeybees, where drone flight times affects mating (Oldroyd et al. 2006), and in the mole salamander, *Ambystoma talpoideum*, where ecological isolation leads to asymmetric reinforcement (Whiteman & Semlitsch, 2005). In this species, paedomorphic and metamorphic individuals interbreed, but RI is stronger in crosses involving metamorphic females, highlighting how spatial and ecological factors drive asymmetric premating isolation (Whiteman & Semlitsch, 2005).

In line with these studies, our findings detected inherently strong mechanical isolation in heterospecific crosses of *I. elegans* androchrome females and *I. graellsii* males, which completely prevents tandem formation. We suggest that this isolation may be linked to the pronotum morphology of androchrome females, potentially reducing excessive conspecific matings. Interestingly, in *I. elegans* conspecific tandem success is similar between male-male and male-androchrome-female interactions (Fig 4). Androchrome females of *I. elegans* are known to mimic males in coloration (Van Gossum et al., 2005), behavior (Cordero-Rivera & Sánchez-Guillén, 2007; Sánchez-Guillén et al., 2017), and abdomen width (Sánchez-Guillén et al., 2005, 2017). It is plausible that they also mimic males in the pronotum, contributing to the higher reproductive isolation observed in heterospecific crosses involving androchrome females.

### Opposite changes between species in the strength of the gametic barriers

Our results showed that changes in the strength of the gametic barriers followed symmetric pattern between morphs, but in opposite directions between species. In *I. elegans* we detected the reinforcement of the fertility barrier, while in *I. graellsii,* we detected the weakening of the fertility and oviposition barriers. Unlike Arce-Valdés et al. (2024), who investigated reinforcement in the NW hybrid zone and detected it only in the mechanical barrier of *I. elegans* females using a limited sample (n = 3, one population pair), our expanded dataset (n = 9, four population pairs) provided greater statistical power to detect gametic reinforcement in this hybrid zone. Consistently with Arce-Valdés et al. (2024) study in the NW hybrid zone we detected the weakening of the fertility barrier in the NC and the NW hybrid zones. In addition, we also detected the weakening in the oviposition barrier but only in the NW hybrid zone. These reductions in fertility and oviposition barriers may be attributed to the high costs of hybridization, which, as shown by previous studies (Arce-Valdés et al., 2024; Ballén-Guapacha et al., 2024), have driven reinforcement and reproductive character displacement in the tandem-related structures of *I. elegans* females and *I. graellsii* males. Furthermore, the ongoing hybridization and bidirectional introgression between these species (Wellenreuther et al., 2018; Swaegers et al., 2022; Sánchez-Guillén et al., 2023) could have facilitated the purging of Bateson-Dobzhansky-Muller incompatibilities, thereby reducing postzygotic isolation in sympatry and leading to the observed weakening of both fertility and oviposition barriers.

Theoretically, reinforcement of gametic barriers can occur even when premating barriers have already evolved, although it is a rare phenomenon with only few documented cases (Johnson & Wade, 1995; Matute, 2010; Turner et al., 2010; Albrecht et al., 2019). The reinforcement of the fertility barrier in our system is consistent with recent models that propose transient fertilization reinforcement can occur when male traits that attract heterospecific females do not reduce conspecific mating success (see Rushworth et al., 2022).

These models predict reinforcement under conditions of recent sympatry, a trade-off in male success, and unidirectional gene flow—conditions that align with our data. Specifically, unidirectional gene flow occurs only between *I. elegans* males and *I. graellsii* females (Sánchez-Guillén et al., 2023), and recent contact is evidenced by the first records of *I. elegans* in the NW hybrid zone, dating back to the early 1980s, and in the NC hybrid zone in the early 2000s.

It remains unclear whether reinforcement of the mechanical and gametic barriers occurred simultaneously or sequentially; however, it has been proposed that, the strengthening of preexisting barriers can drive the emergence or further strengthening of other barriers through a coupling effect (Butlin & Smadja, 2018), which can occur through mechanisms such as spatial shifts in clines (Bierne et al., 2011), population dynamics like extinction-recolonization, or brief periods of allopatry where locally adaptive loci also contribute to hybrid incompatibilities (Kulmuni et al., 2020). Given that extinction-recolonization events are common in the two hybrid zones we studied (Sánchez-Guillén et al., 2023), it is plausible that the reinforcement of mechanical barriers facilitated the reinforcement of gametic barriers through a coupling effect. Future research should assess linkage disequilibrium between barrier loci to evaluate the strength of coupling and the relative roles of secondary contact and reinforcement.

If these barriers were reinforced sequentially, their relative contribution to reproductive isolation in sympatry may have shifted rapidly over time (e.g. Castillo & Moyle, 2019). However, the fact that two barriers have been reinforced in this system raises an interesting question: why does selection reinforce two barriers instead of completing speciation through a single barrier? One possibility is that reinforcement of the mechanical barrier was still incomplete at the time of sampling and could eventually lead to full speciation. Alternatively, further reinforcement of the mechanical barrier may have been constrained by the coevolution of male caudal appendages and female prothorax in a lock-and-key pattern (Ballén-Guapacha et al., 2024). If selection intensified mechanical isolation, it could disrupt tandem formation in conspecifics, as male structures might not evolve quickly enough to match female modifications. This is supported by phylogenetic studies showing that female genitalia evolve up to three times faster than male genitalia in some insects (Simmons & Fitzpatrick, 2019), potentially limiting the extent of mechanical reinforcement.

### Evolutionary consequences of the asymmetric reinforcement

As reinforcement strengthens reproductive barriers in only one of the morphs—as observed in *I. elegans* gynochromes—it may trigger long-term evolutionary consequences. The reinforced morph could experience greater reproductive isolation, diverging from both the ancestral population and the non-reinforced morph (West-Eberhard, 1986; Corl et al., 2010; Hugall & Stuart-Fox, 2012; Heuer et al., 2024). Evidence for morph-driven speciation exists in the lizard *Uta stansburiana*, where polymorphism has persisted for millions of years, yet populations that lost morphs underwent rapid phenotypic evolution and subspeciation (Corl et al., 2010). Unlike West-Eberhard’s (1986) classic hypothesis, where morph loss or gain drives speciation, we propose that in our case, reproductive isolation evolves first in sympatry, acting as a driver rather than a consequence of speciation.

## Conclusions

Our findings underscore the complex nature of the evolutionary process during hybridization between *I. graellsii* and *I. elegans*. The asymmetrical pattern of reinforcement between female morphs highlights the differential selection pressures acting on androchrome and gynochrome females, influencing their reproductive strategies and hybridization dynamics. Here, we provide insights into the early stages of speciation and emphasize the role of RI in maintaining species boundaries and promoting biodiversity. Moreover, our findings provide a clear example of asymmetrical reinforcement between morphs, illustrating how polymorphism can increase the likelihood of reinforcement due to inherent asymmetries in RI between morphs and allowed us to propose an alternative hypothesis in which morph speciation can take place. Therefore, our work sheds light on the complex evolutionary dynamic between polymorphism and reinforcement. Future work should focus on exploring how asymmetric reinforcement influence the evolution and maintenance of polymorphism, shaping morph frequencies in sympatry.

## AUTHOR CONTRIBUTIONS

**Conceptualization:** J.E.O.-M. and R.A.S.-G., **Data curation:** J.E.O.-M., A.L.J.-J., M.S.-P., L.R.A.-V., A.V.B.-G., J.R.C.-R. and R.A.S.-G., **Formal analysis:** J.E.O.-M., L.R.A.-V., and R.A.S.-G., **Visualization:** J.E.O.-M. and R.A.S.-G., **Writing-original draft:** J.E.O.-M. and R.A.S.-G., **Writing—review & editing:** all the authors, **Funding acquisition: A.C.-R.** and R.A.S.-G.

## ACKNOWLEDGEMENTS

We would like to thank the Zalandrana Odonatology group who kindly helped us with sampling and permitting in North-central Spain, to Janet Nolasco Soto and Jovita Martínez Lapa for their technical support during damselfly rearing, to Samantha Maíte de los Santos Gómez for her assistance in map preparation, Roger Guevara for his statistical advice and Angela Nava Bolaños and Oscar Rios-Cardenas for comments on the first draft of the manuscript. J.E.O.-M utilized language processing tools driven by artificial intelligence (i.e., ChatGPT) to check and/or fix syntax during R code writing and grammar correction on the main text.

## Supplementary materials

**Table S1.**
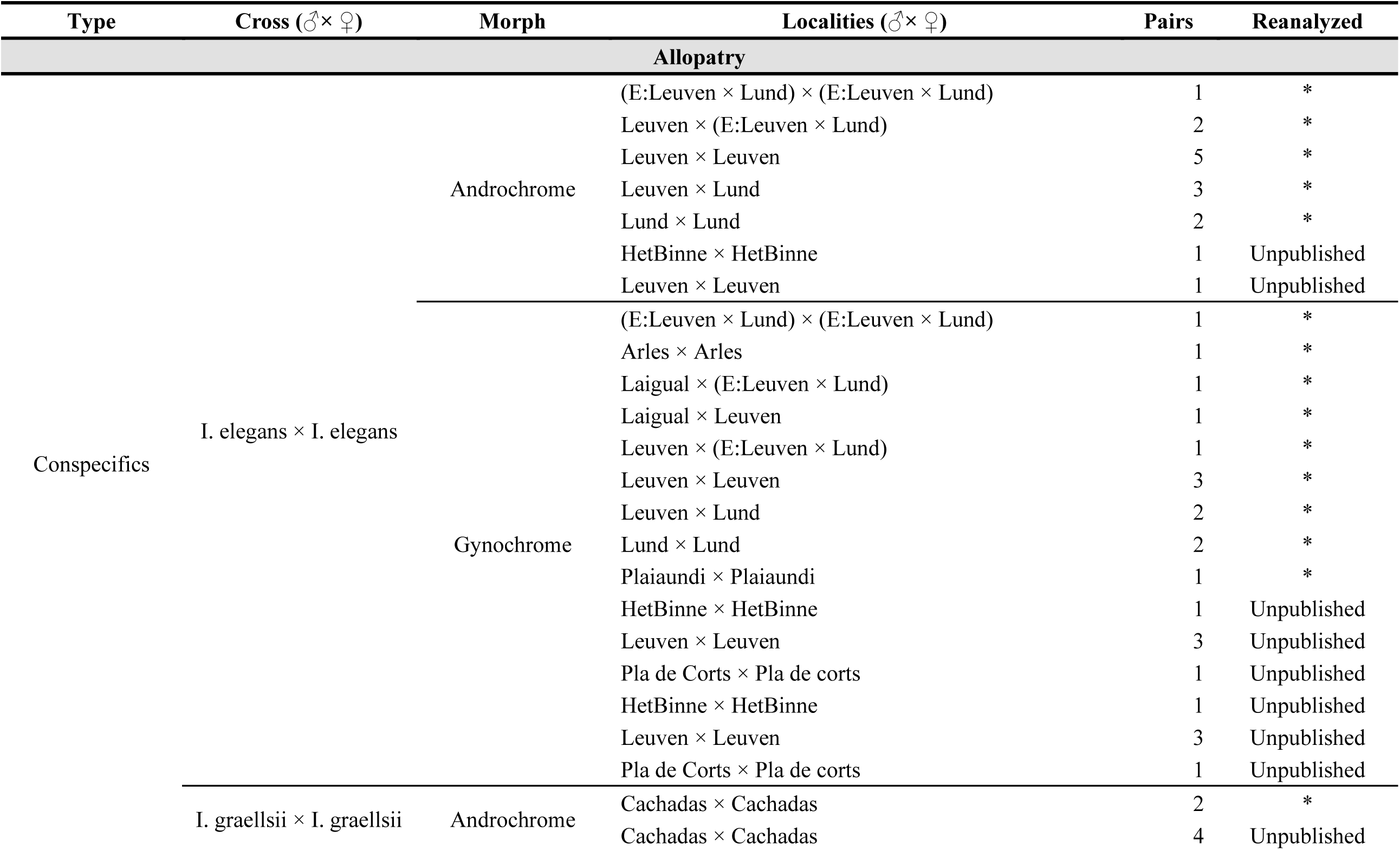
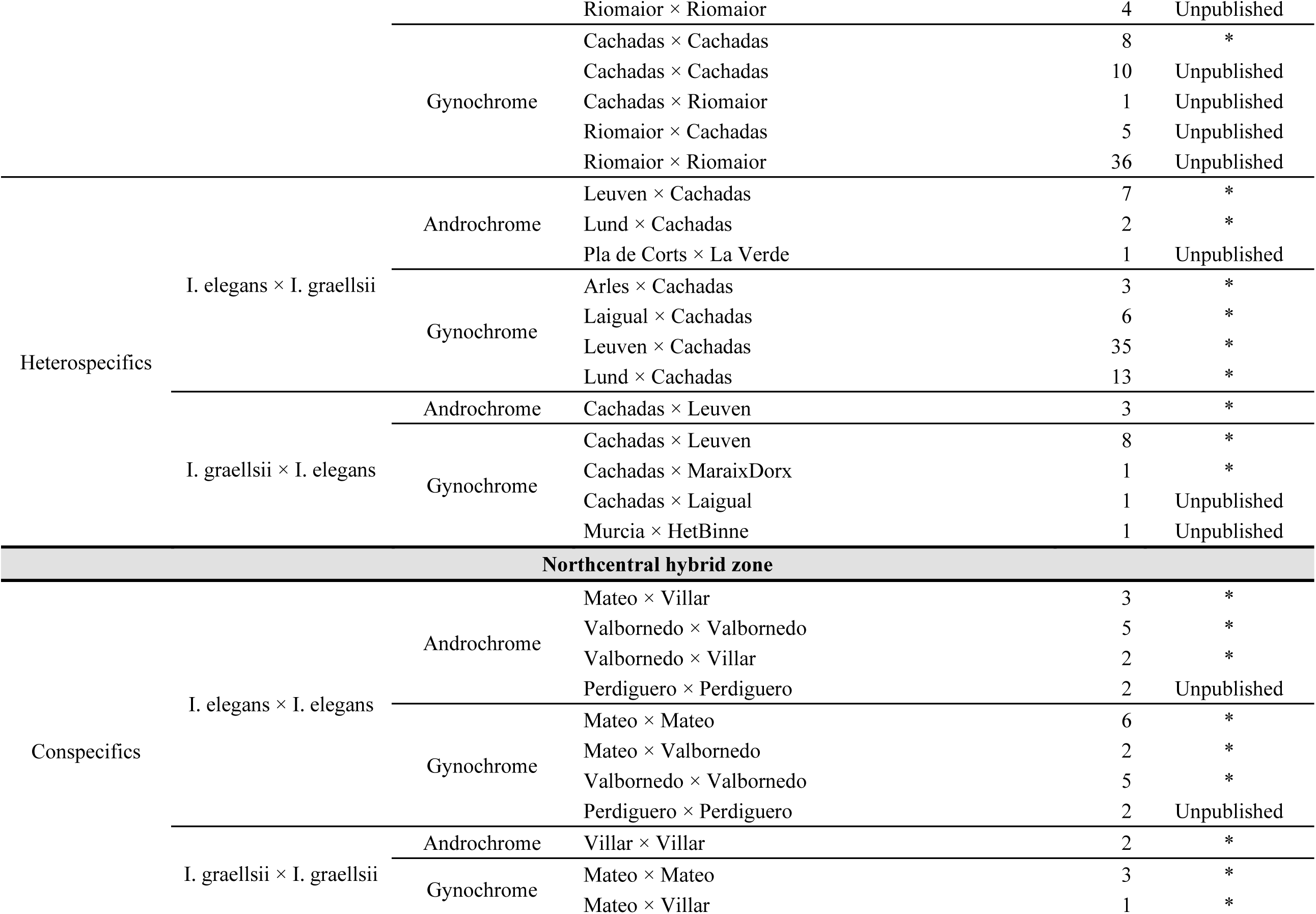
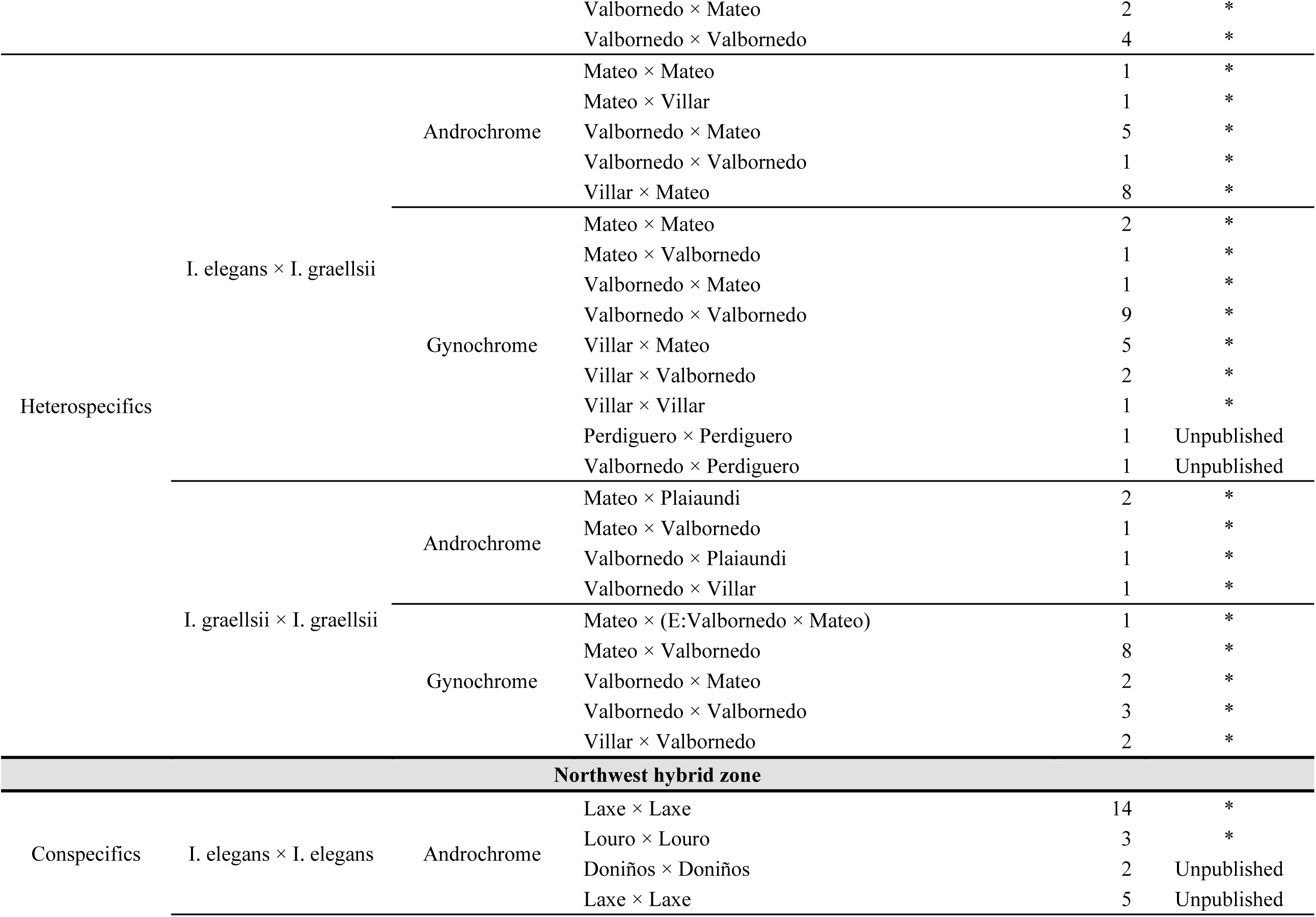
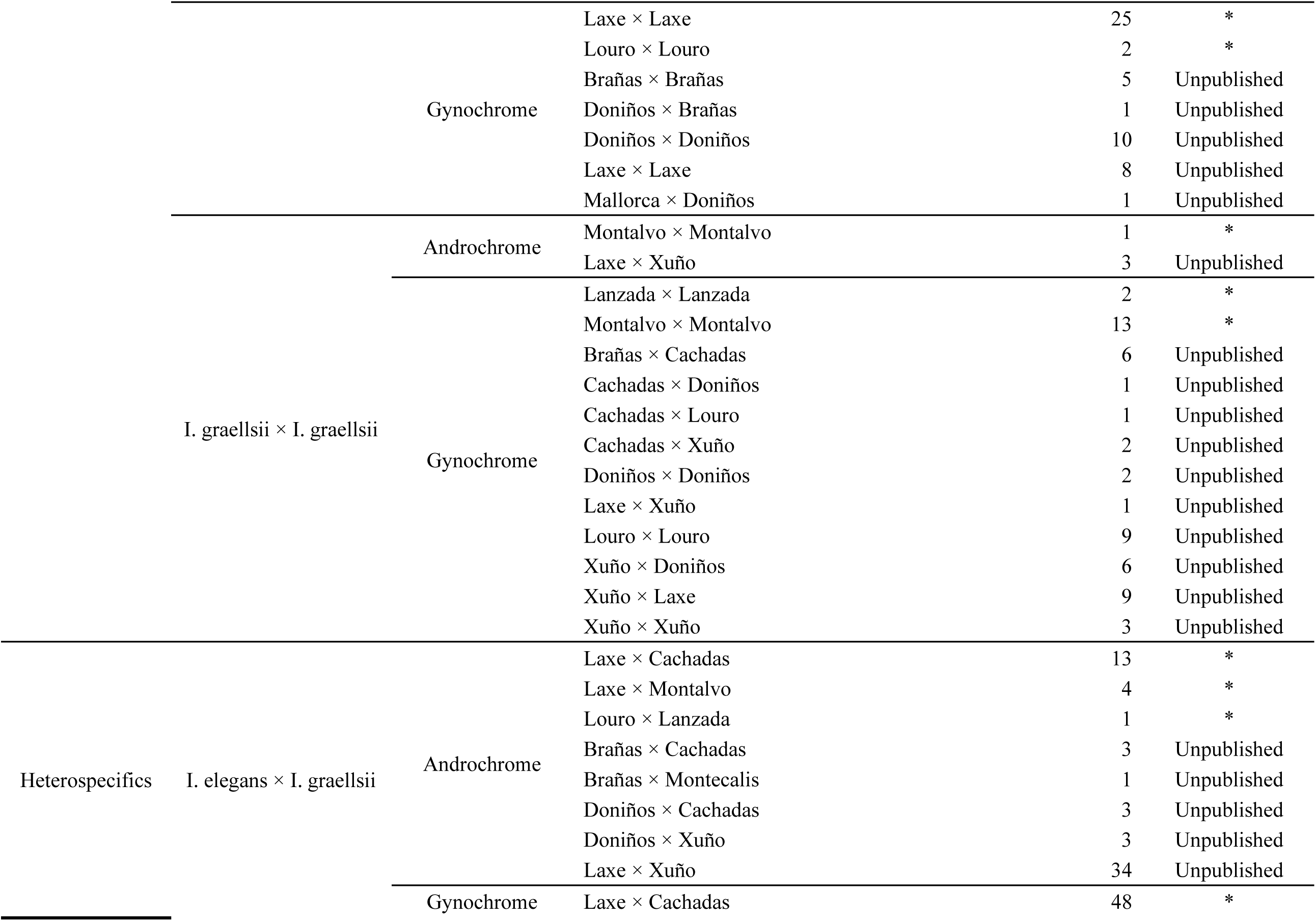
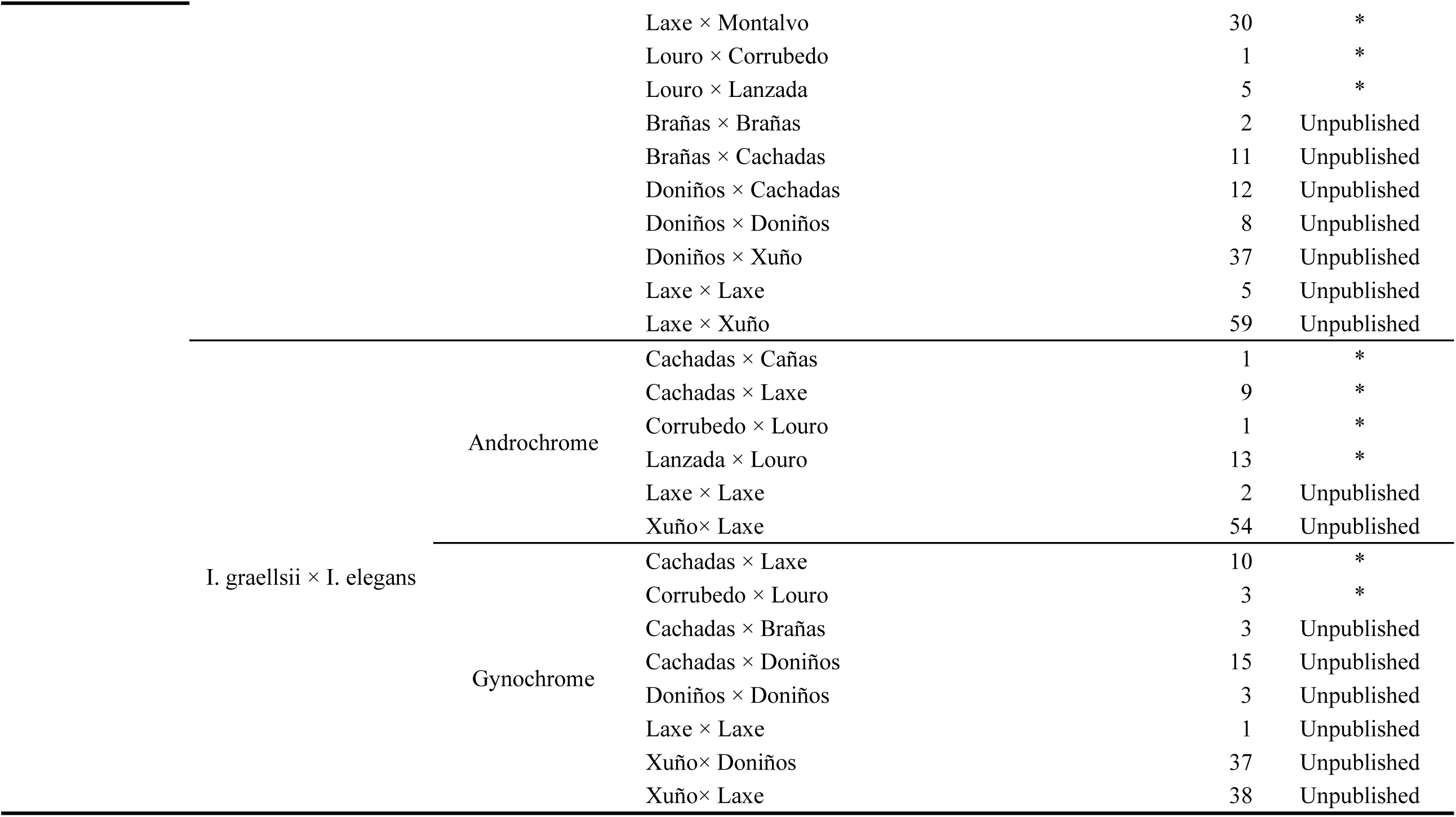
Number of unique mating pairs from which the data of five prezygotic reproductive barriers were collected. Reproductive isolation between conspecifics was measured to be used as controls for heterospecific crosses. The asterisk in last column indicated data reanalyzed from Sánchez-Guillén et al. (2012), Sánchez-Guillén et al. (2023) and Arce-Valdes et al. (2024).

**Table S2.** Mating success rates in conspecific crosses, used as control values, for the five measured reproductive barriers: Mechanical— measured as the rate of successful tandems over tandem attempts; Mechanical-Tactile—measured as the rate of successful matings over tandems; Oviposition—measured as the proportion of mated females that successfully laid eggs; Fecundity—measured as the average number of laid egg per female during the first three ovipositions; and Fertility—measured as the proportion of fertile eggs by female. The average success rate (Avg success) is the mean conspecific success of the given female morph between both species. This is the value we used as control in our estimation of heterospecific reproductive isolation (formula 1 in main document). The average isolation (Avg isolation) is the complement of the average success rate, indicating the average conspecific failure of a given female morph between both species, estimated as 1 – Avg success.

| Barrier | Androchrome morph |  |  |  | Gynochrome morph |  |  |  |
| --- | --- | --- | --- | --- | --- | --- | --- | --- |
|  | *I. elegans* | *I. graellsii* | Avg success | Avg isolation | *I. elegans* | *I. graellsii* | Avg success | Avg Isolation |
| Allopatry |  |  |  |  |  |  |  |  |
| Mechanical | 0.895 | 0.435 | 0.665 | 0.335 | 0.682 | 0.466 | 0.574 | 0.426 |
| Mechanical-Tactile | 0.824 | 0.500 | 0.662 | 0.338 | 0.933 | 0.869 | 0.901 | 0.099 |
| Oviposition | 0.895 | 0.917 | 0.906 | 0.094 | 0.923 | 1.000 | 0.962 | 0.038 |
| Fecundity | 211.010 | 168.265 | 189.638 | 189.638 | 236.856 | 170.579 | 203.718 | 203.718 |
| Fertility | 0.609 | 0.798 | 0.704 | 0.297 | 0.698 | 0.872 | 0.785 | 0.215 |
| Northcentral hybrid zone |  |  |  |  |  |  |  |  |
| Mechanical | 0.583 | 0.600 | 0.592 | 0.408 | 0.529 | 0.900 | 0.715 | 0.285 |
| Mechanical-Tactile | 0.857 | 0.667 | 0.762 | 0.238 | 0.444 | 1.000 | 0.722 | 0.278 |
| Oviposition | 0.889 | 1.000 | 0.944 | 0.056 | 0.909 | 1.000 | 0.955 | 0.045 |
| Fecundity | 124.458 | 70.333 | 97.396 | 97.396 | 198.267 | 128.653 | 163.460 | 163.460 |
| Fertility | 0.439 | 0.319 | 0.379 | 0.621 | 0.575 | 0.435 | 0.505 | 0.495 |
| Northwest hybrid zone |  |  |  |  |  |  |  |  |
| Mechanical | 0.484 | 0.500 | 0.492 | 0.508 | 0.595 | 0.586 | 0.591 | 0.409 |
| Mechanical-Tactile | 0.600 | 0.500 | 0.550 | 0.450 | 0.660 | 0.686 | 0.673 | 0.327 |
| Oviposition | 0.966 | 1.000 | 0.983 | 0.017 | 0.944 | 0.977 | 0.961 | 0.039 |
| Fecundity | 171.757 | 222.000 | 196.879 | 196.879 | 188.162 | 282.324 | 235.243 | 235.243 |
| Fertility | 0.807 | 0.970 | 0.889 | 0.112 | 0.842 | 0.981 | 0.912 | 0.089 |

**Table S3.** Cumulative total reproductive isolation per barrier in heterospecific crosses, measured in allopatry, hybrid zones and sympatry (both hybrid zones together). Bold values indicate complete reproductive isolation, while negative values indicate facilitated hybridization. Sample size refers to the number of mating pairs. E × G represents *I. elegans* x *I. graellsii*, G × E represents *I. graellsii* x *I. elegans*. A denotes androchrome females, and G gynochrome females.

| Cross (♂×♀) | Morph | Sample size | Mechanical | Mechanical-Tactile | Oviposition | Fecundity | Fertility |
| --- | --- | --- | --- | --- | --- | --- | --- |
| **Allopatry** |  |  |  |  |  |  |  |
| E × G | A | 10 | 0.074 | -0.087 | -0.200 | 0.887 | **1.000** |
|  | G | 57 | -0.044 | 0.063 | 0.338 | 0.812 | 0.988 |
| G × E | A | 3 | 0.614 | 0.483 | 0.429 | 0.119 | -0.012 |
|  | G | 11 | -0.266 | -0.149 | -0.264 | -0.872 | -0.748 |
| **Northcentral hybrid region** |  |  |  |  |  |  |  |
| E × G | A | 16 | 0.053 | 0.320 | 0.280 | -1.201 | -1.874 |
|  | G | 23 | 0.130 | 0.141 | 0.164 | 0.733 | 0.912 |
| G × E | A | 5 | 0.495 | 0.456 | 0.424 | -0.433 | 0.853 |
|  | G | 16 | 0.632 | 0.647 | 0.630 | 0.817 | 0.778 |
| **Northwest hybrid region** |  |  |  |  |  |  |  |
| E × G | A | 62 | 0.097 | 0.097 | 0.132 | 0.480 | 0.535 |
|  | G | 218 | 0.220 | 0.304 | 0.326 | 0.598 | 0.698 |
| G × E | A | 80 | 0.940 | 0.965 | 0.964 | 0.972 | 0.978 |
|  | G | 110 | 0.868 | 0.910 | 0.906 | 0.954 | 0.966 |
| **Sympatry** |  |  |  |  |  |  |  |
| E × G | A | 73 | 0.206 | 0.348 | 0.373 | 0.398 | 0.288 |
|  | G | 207 | 0.265 | 0.352 | 0.386 | 0.616 | 0.718 |
| G × E | A | 72 | 0.923 | 0.924 | 0.922 | 0.892 | 0.955 |
|  | G | 126 | 0.839 | 0.875 | 0.870 | 0.934 | 0.947 |

**Table S4.** *Post hoc* contrast tests modeling for reproductive isolation as a function of the interaction between types of crosses, distribution and morph (RI ∼ interaction) per prezygotic reproductive barrier using the hybrid zone and regional dataset to detect statistical differences between allopatric and each hybrid zone, and sympatry, per morph. GLMs were fitted using each morph as model intercept to allow pairwise comparisons between morphs. Bold values indicate significant *p* value for differences between a morph and the model intercept following the sequential Bonferroni correction (*p* < 0.05/6 and *p* < 0.05/4 for the hybrid zone and regional data set, respectively). **Note: Contrast tests were not performed at local level due to the low sample size for most of the localities.**

| Cross (♂ × ♀) | Pairwise | Morph | Odds ratio / estimate | S.E. | df | z or t value | *P* adj | Holm threshold |
| --- | --- | --- | --- | --- | --- | --- | --- | --- |
| Hybrid zone data set |  |  |  |  |  |  |  |  |
| *Premating I – Mechanical barrier (Binomial distribution)* |  |  |  |  |  |  |  |  |
| I. elegans × I. graellsii | Allopatry VS NCHZ | Androchrome | 0.7617 | 0.4171 | - | -0.4971 | 1.0000 | 0.0292 |
|  |  | Gynochrome | 0.6914 | 0.2440 | - | -1.0455 | 1.0000 | 0.0417 |
|  | Allopatry VS NWHZ | Androchrome | 0.5030 | 0.2362 | - | -1.4632 | 1.0000 | 0.0333 |
|  |  | Gynochrome | 0.4794 | 0.1041 | - | -3.3869 | 0.0085 | 0.0083 |
|  | NCHZ VS NWHZ | Androchrome | 0.6604 | 0.2444 | - | -1.1209 | 1.0000 | 0.0375 |
|  |  | Gynochrome | 0.6934 | 0.2150 | - | -1.1808 | 1.0000 | 0.0458 |
| I. graellsii × I. elegans | Allopatry VS NCHZ | Androchrome | 1.6000 | 2.2343 | - | 0.3366 | 1.0000 | 0.0500 |
|  |  | Gynochrome | 0.2174 | 0.1392 | - | -2.3829 | **0.0206** | 0.0208 |
|  | Allopatry VS NWHZ | Androchrome | 0.0678 | 0.0855 | - | -2.1350 | 0.3931 | 0.0250 |
|  |  | Gynochrome | 0.0313 | 0.0160 | - | -6.7857 | **< 0.0001** | 0.0042 |
|  | NCHZ VS NWHZ | Androchrome | 0.0424 | 0.0432 | - | -3.1014 | 0.0231 | 0.0167 |
|  |  | Gynochrome | 0.1442 | 0.0837 | - | -3.3352 | **0.0102** | 0.0125 |
| *Premating II – Mechanical-tactile barrier (Binomial distribution)* |  |  |  |  |  |  |  |  |
| I. elegans × I. graellsii | Allopatry VS NCHZ | Androchrome | 0.7033 | 0.4820 | - | -0.5135 | 1.0000 | 0.0042 |
|  |  | Gynochrome | 0.9825 | 0.4100 | - | -0.0424 | 1.0000 | 0.0167 |
|  | Allopatry VS NWHZ | Androchrome | 0.5455 | 0.3172 | - | -1.0423 | 1.0000 | 0.0083 |
|  |  | Gynochrome | 0.8395 | 0.2102 | - | -0.6987 | 1.0000 | 0.0208 |
|  | NCHZ VS NWHZ | Androchrome | 0.7756 | 0.4047 | - | -0.4871 | 1.0000 | 0.0125 |
|  |  | Gynochrome | 0.8545 | 0.3226 | - | -0.4165 | 1.0000 | 0.0250 |
| I. graellsii × I. elegans | Allopatry VS NCHZ | Androchrome | 1.0000 | 1.7321 | - | 0.0000 | 1.0000 | 0.0292 |
|  |  | Gynochrome | 1.0667 | 0.8563 | - | 0.0804 | 1.0000 | 0.0417 |
|  | Allopatry VS NWHZ | Androchrome | 0.3333 | 0.6086 | - | -0.6017 | 1.0000 | 0.0333 |
|  |  | Gynochrome | 0.7273 | 0.4890 | - | -0.4737 | 1.0000 | 0.0458 |
|  | NCHZ VS NWHZ | Androchrome | 0.3333 | 0.5092 | - | -0.7192 | 1.0000 | 0.0375 |
|  |  | Gynochrome | 0.6818 | 0.5735 | - | -0.4553 | 1.0000 | 0.0500 |
| Postmating I – Oviposition (Binomial distribution) |  |  |  |  |  |  |  |  |
| I. elegans × I. graellsii | Allopatry VS NCHZ | Androchrome | 1.0000 | 5325.7119 | - | 0.0000 | 1.0000 | 0.0125 |
|  |  | Gynochrome | 0.1385 | 0.1537 | - | -1.7810 | 0.8990 | 0.0083 |
| | Allopatry VS NWHZ | Androchrome | $6.8028 \times 10^6$ | $2.5618 \times 10^{10}$ | - | 0.0042 | 1.0000 | 0.0167 |
|  |  | Gynochrome | 0.1355 | 0.0753 | - | -3.5950 | **0.0039** | 0.0042 |
| | NCHZ VS NWHZ | Androchrome | $6.8028 \times 10^6$ | $2.5618 \times 10^{10}$ | - | 0.0042 | 1.0000 | 0.0208 |
|  |  | Gynochrome | 0.9785 | 1.0854 | - | -0.0196 | 1.0000 | 0.0250 |
| I. graellsii × I. elegans | Allopatry VS NCHZ | Androchrome | 1.0000 | 7988.5682 | - | 0.0000 | 1.0000 | 0.0292 |
|  |  | Gynochrome | 1.0000 | 4210.3451 | - | 0.0000 | 1.0000 | 0.0417 |
|  | Allopatry VS NWHZ | Androchrome | 1.0000 | 7988.5681 | - | 0.0000 | 1.0000 | 0.0333 |
|  |  | Gynochrome | 1.0000 | 3628.8611 | - | 0.0000 | 1.0000 | 0.0458 |
|  | NCHZ VS NWHZ | Androchrome | 1.0000 | 6522.6386 | - | 0.0000 | 1.0000 | 0.0375 |
|  |  | Gynochrome | 1.0000 | 4088.2801 | - | 0.0000 | 1.0000 | 0.0500 |
| Postmating II – Gametic (Fecundity) (Gamma distribution) |  |  |  |  |  |  |  |  |
| I. elegans × I. graellsii | Allopatry VS NCHZ | Androchrome | -199.3333 | 122.3451 | 156 | -1.6293 | 1.0000 | 0.0208 |
|  |  | Gynochrome | 7.3889 | 54.5387 | 156 | 0.1355 | 1.0000 | 0.0333 |
|  | Allopatry VS NWHZ | Androchrome | 191.5000 | 183.5176 | 156 | 1.0435 | 1.0000 | 0.0417 |
|  |  | Gynochrome | -120.1329 | 38.6188 | 156 | -3.1107 | 0.0266 | 0.0042 |
|  | NCHZ VS NWHZ | Androchrome | 82.1765 | 93.8344 | 156 | 0.8758 | 1.0000 | 0.0292 |
|  |  | Gynochrome | -127.5217 | 44.3978 | 156 | -2.8723 | 0.0557 | 0.0125 |
| I. graellsii × I. elegans | Allopatry VS NCHZ | Androchrome | 16.5000 | 183.5176 | 156 | 0.0899 | 1.0000 | 0.0375 |
|  |  | Gynochrome | 252.1667 | 96.7223 | 156 | 2.6071 | 0.1202 | 0.0167 |
|  | Allopatry VS NWHZ | Androchrome | 191.5000 | 183.5176 | 156 | 1.0435 | 1.0000 | 0.0417 |
|  |  | Gynochrome | 257.381 | 83.3641 | 156 | 3.0874 | 0.0287 | 0.0083 |
|  | NCHZ VS NWHZ | Androchrome | 175.0000 | 149.8415 | 156 | 1.1679 | 1.0000 | 0.0458 |
|  |  | Gynochrome | 5.2143 | 93.9181 | 156 | 0.0555 | 1.0000 | 0.0500 |

| Postmating III – Gametic (Fertility) (Binomial distribution) |  |  |  |  |  |  |  |  |
| --- | --- | --- | --- | --- | --- | --- | --- | --- |
| I. elegans × I. graellsii | Allopatry VS NCHZ | Androchrome | 0.0190 | 0.0191 | - | -3.9403 | **0.0010** | 0.0458 |
|  |  | Gynochrome | 2.1979 | 0.1821 | - | 9.5047 | **< 0.0001** | 0.0292 |
|  | Allopatry VS NWHZ | Androchrome | 0.0015 | 0.0015 | - | -6.4836 | **< 0.0001** | 0.0375 |
|  |  | Gynochrome | 0.2613 | 0.0139 | - | -25.1829 | **< 0.0001** | 0.0167 |
|  | NCHZ VS NWHZ | Androchrome | 0.0782 | 0.0063 | - | -31.8782 | **< 0.0001** | 0.0042 |
|  |  | Gynochrome | 0.1189 | 0.0080 | - | -31.4580 | **< 0.0001** | 0.0083 |
| I. graellsii × I. elegans | Allopatry VS NCHZ | Androchrome | 74.6920 | 12.3878 | - | 26.0074 | **< 0.0001** | 0.0125 |
|  |  | Gynochrome | 3.0360 | 0.2252 | - | 14.9716 | **< 0.0001** | 0.0250 |
|  | Allopatry VS NWHZ | Androchrome | 1.8658 | 0.3760 | - | 3.0949 | **0.0236** | 0.0500 |
|  |  | Gynochrome | 1.7944 | 0.1207 | - | 8.6920 | **< 0.0001** | 0.0333 |
|  | NCHZ VS NWHZ | Androchrome | 0.0250 | 0.0055 | - | -16.7858 | **< 0.0001** | 0.0208 |
|  |  | Gynochrome | 0.5910 | 0.0503 | - | -6.1754 | **< 0.0001** | 0.0417 |
| Regional dataset |  |  |  |  |  |  |  |  |
| Premating I – Mechanical barrier (Binomial distribution) |  |  |  |  |  |  |  |  |
| I. elegans × I. graellsii | - | Androchrome | 0.5040 | 0.2348 | - | -1.4706 | 0.5656 | 0.0500 |
|  | - | Gynochrome | 0.4622 | 0.1010 | - | -3.5317 | **0.0017** | 0.0250 |
| I. graellsii × I. elegans | - | Androchrome | 0.1006 | 0.1235 | - | -1.8710 | 0.2454 | 0.0375 |
|  | - | Gynochrome | 0.0428 | 0.0206 | - | -6.5455 | **< 0.0001** | 0.0125 |
| Premating II – Mechanical-tactile barrier (Binomial distribution) |  |  |  |  |  |  |  |  |
| I. elegans × I. graellsii | - | Androchrome | 0.6171 | 0.3518 | - | -0.8468 | 1.0000 | 0.0125 |
|  | - | Gynochrome | 0.8253 | 0.2088 | - | -0.759 | 1.0000 | 0.0250 |
| I. graellsii × I. elegans | - | Androchrome | 0.75 | 1.2055 | - | -0.179 | 1.0000 | 0.0375 |
|  | - | Gynochrome | 0.8333 | 0.4979 | - | -0.3052 | 1.0000 | 0.0500 |
| Postmating I – Oviposition (Binomial distribution) |  |  |  |  |  |  |  |  |
| I. elegans × I. graellsii | - | Androchrome | $7.7099 \times 10^6$ | $2.9034 \times 10^{10}$ | - | 0.0042 | 1.0000 | 0.0250 |
|  | - | Gynochrome | 0.1756 | 0.0950 | - | -3.2149 | **0.0052** | 0.0125 |
| I. graellsii × I. elegans | - | Androchrome | 1.0000 | 7292.5316 | - | 0.0000 | 1.0000 | 0.0375 |
|  | - | Gynochrome | 1.0000 | 3310.3644 | - | 0.0000 | 1.0000 | 0.0500 |
| Postmating II – Gametic (Fecundity) (Gamma distribution) |  |  |  |  |  |  |  |  |
| I. elegans × I. graellsii | - | Androchrome | -110.6000 | 92.3221 | 131 | -1.198 | 0.9324 | 0.0375 |
|  | - | Gynochrome | -98.3765 | 38.0377 | 131 | -2.5863 | 0.0432 | 0.0250 |
| I. graellsii × I. elegans | - | Androchrome | 104.0000 | 163.2039 | 131 | 0.6372 | 1.0000 | 0.0500 |
|  | - | Gynochrome | 255.4848 | 74.0846 | 131 | 3.4486 | **0.0030** | 0.0125 |
| *Postmating III – Gametic (Fertility) (Binomial distribution)* |  |  |  |  |  |  |  |  |
| I. elegans × I. graellsii | - | Androchrome | 0.0026 | 0.0026 | - | -5.9472 | < **0.0001** | 0.0500 |
|  | - | Gynochrome | 0.3461 | 0.0184 | - | -19.9166 | < **0.0001** | 0.0250 |
| I. graellsii × I. elegans | - | Androchrome | 28.1885 | 3.6563 | - | 25.7415 | < **0.0001** | 0.0125 |
|  | - | Gynochrome | 2.2377 | 0.1254 | - | 14.3762 | < **0.0001** | 0.0375 |

**Table S5.**
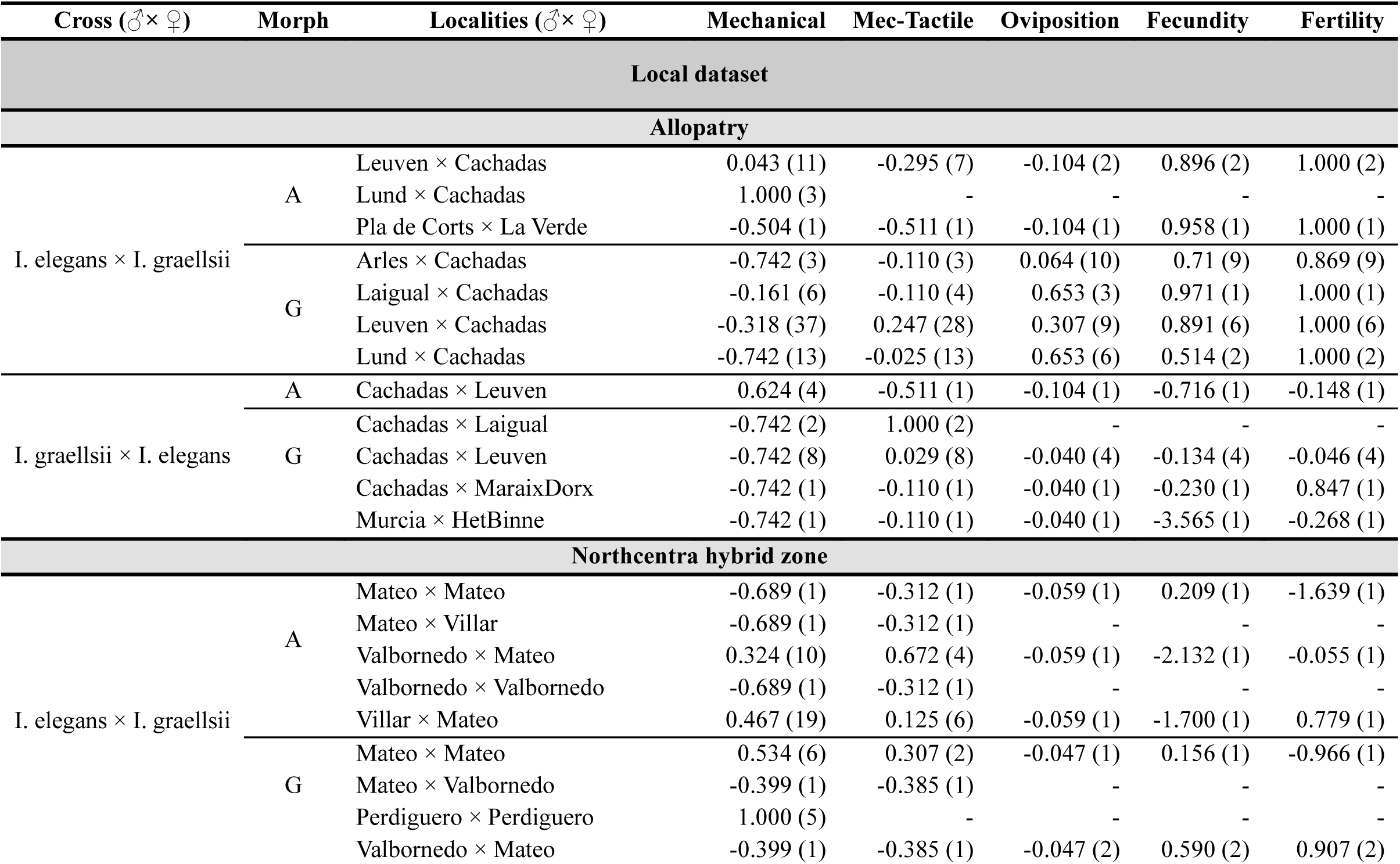
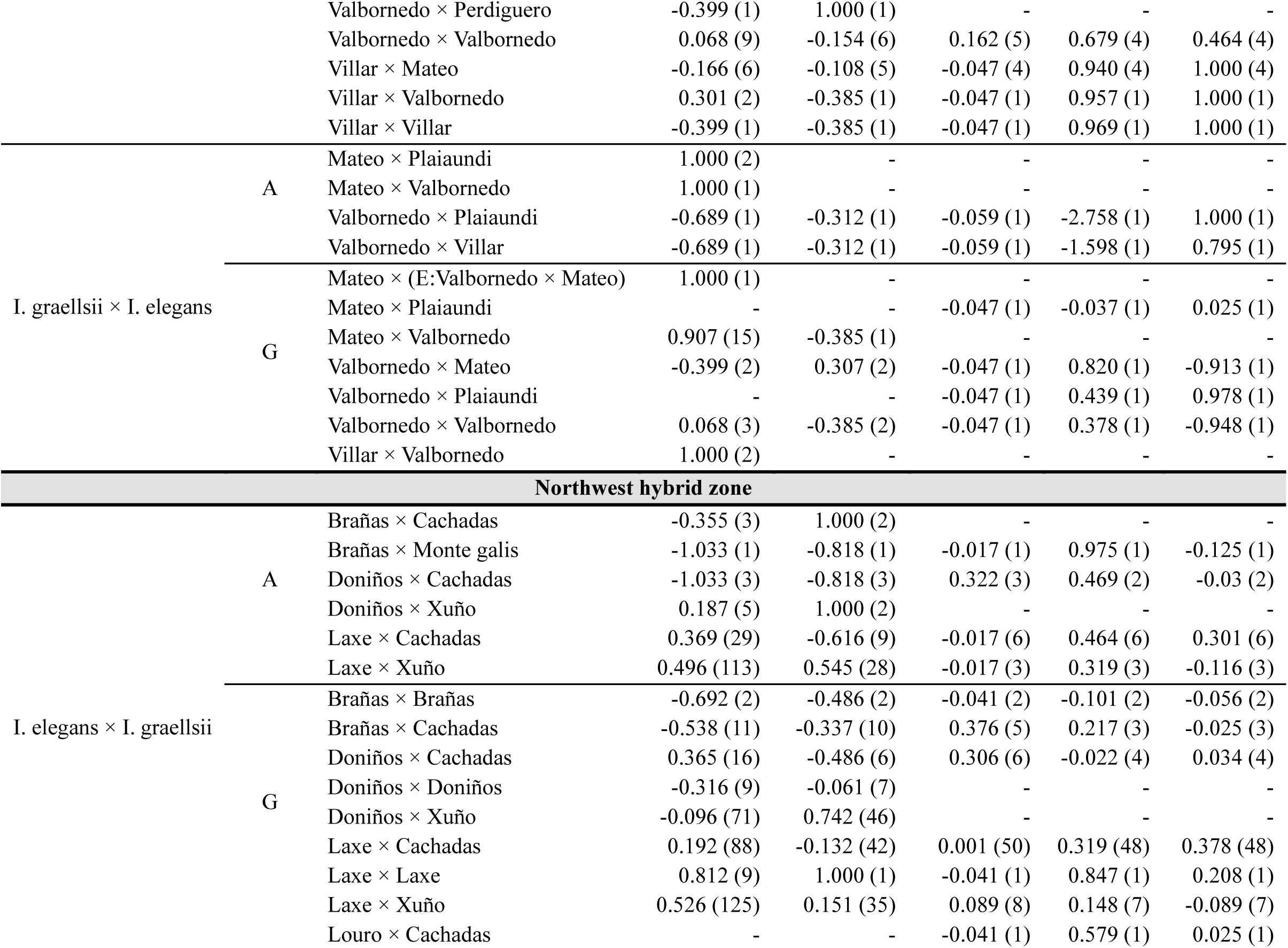
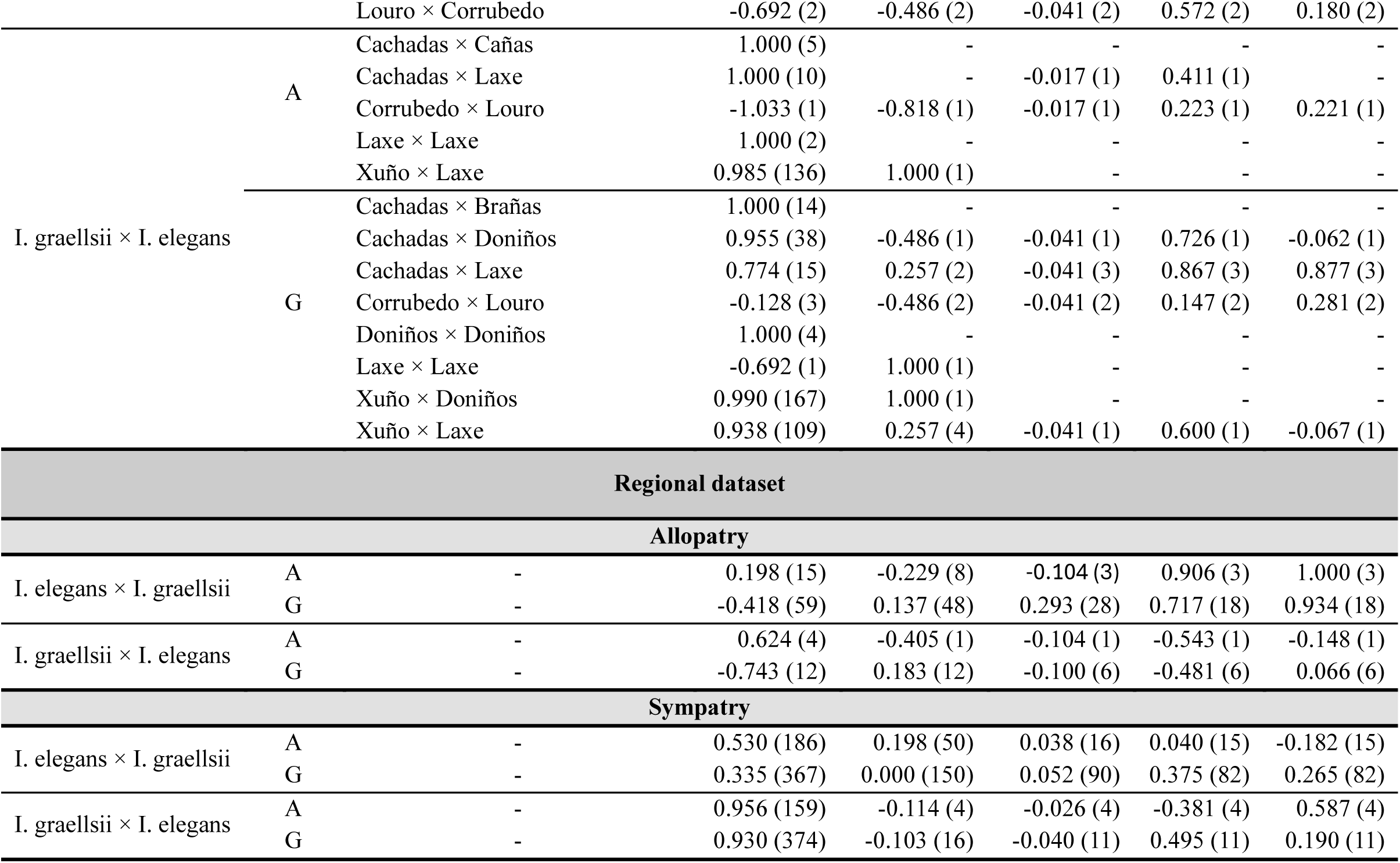
Absolute reproductive isolation and sample size (between parentheses) per reproductive barrier measured for heterospecific crosses at population level (localities) in allopatry, Northwest and Northcentral hybrid zone. Absolute reproductive isolation and sample size are also presented for the regional dataset (allopatry vs sympatry). Sample size in mechanical and mechanical-tactile barriers refers to the number of tandem attempts and tandems, respectively, performed by each cross direction. Bootstrap analysis to estimate 95% CI was not performed for the local data set due to multiple crosses with NA and/or n = 1 for the measured reproductive barriers.

**Figure S1.**
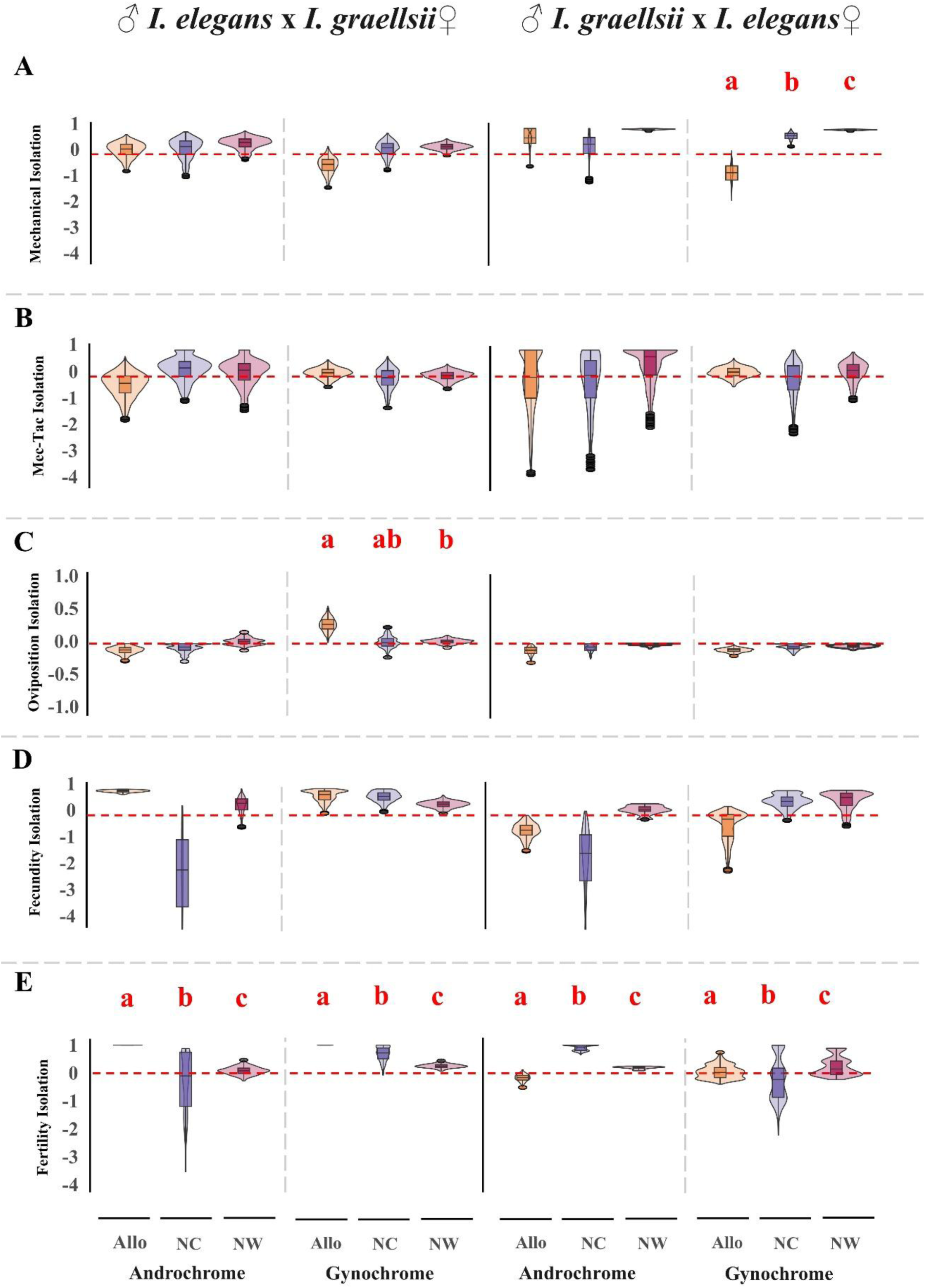
Absolute strength of each reproductive barrier and 95%CI between hybrid zones. **A)** mechanical isolation measured as the number of tandem attempts per heterospecific pair. **B)** mechanical-tactile isolation, measured as the number of mating attempts per heterospecific pair. **C)** oviposition isolation, measured as the number of females that oviposited after mating. **D)** gametic isolation, fecundity, measured as the number of eggs laid by each female. **E)** gametic isolation, fertility, measured as the ratio of fertile eggs laid by each female. Each barrier is presented with 95%CI estimated from 10,000 bootstrap resample of the dataset. The horizontal red dashed line marks the boundary between reproductive isolation (above the line) and facilitated hybridization (below the line). Red bold letters indicate statistically significant difference between the allopatric (boxplot and violin), NC hybrid zone (purple boxplot and violin), and NW hybrid zone (red boxplot and violin) after Holm-Bonferroni correction. Analysis was performed with the raw data.

**Figure S2.**
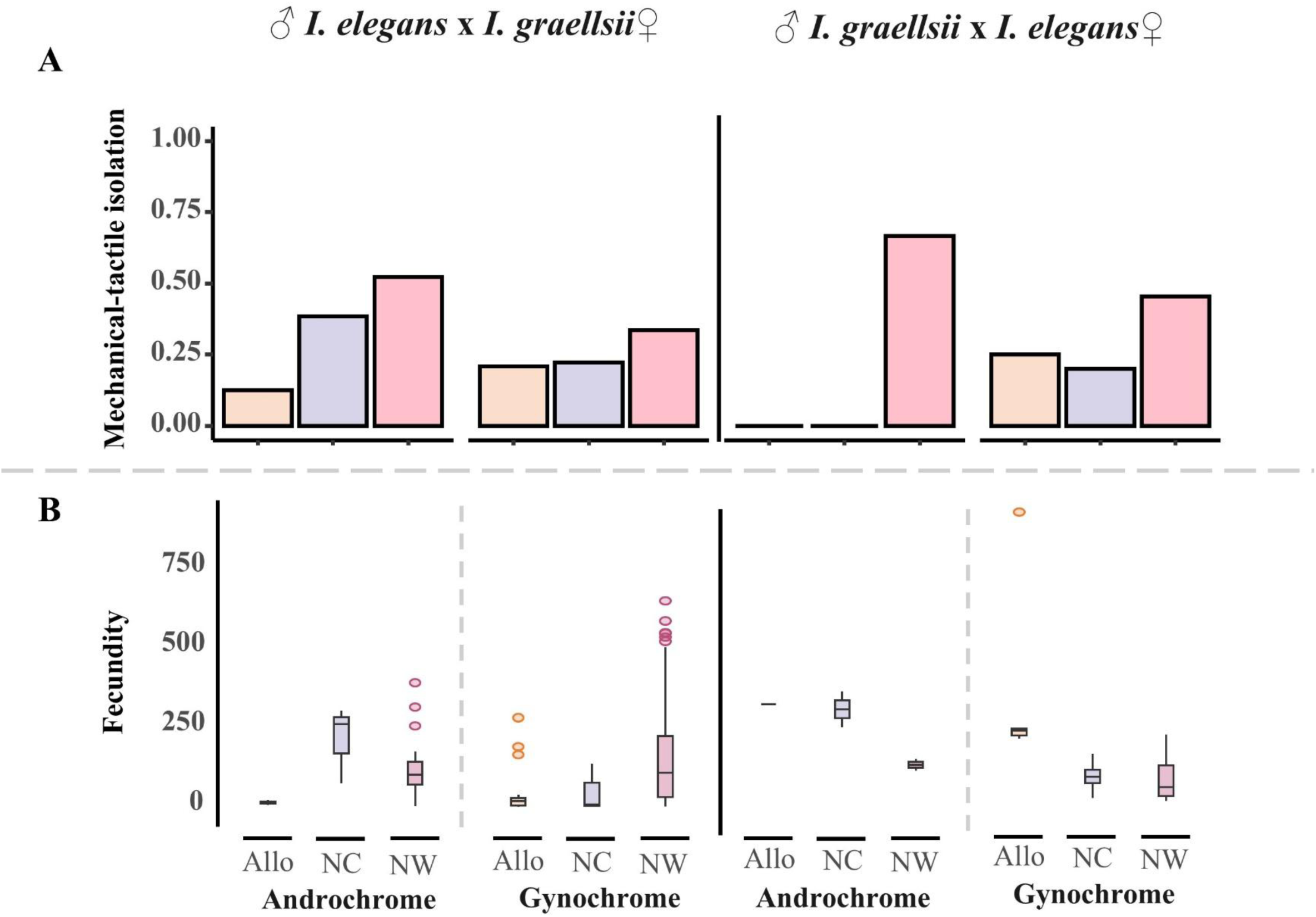
Differences in mechanical-tactile barrier and fecundity (gametic barrier) between hybrid zones. **A)** Mechanical-tactile isolation, measured as the number of mating attempts per heterospecific pair. **B)** fecundity, measured as the number of eggs laid by each female. No statistically significant difference was found between the allopatric (orange barplot and boxplot), NC hybrid zone (purple barplot and boxplot), and NW hybrid zone (pink barplot and boxplot) after Holm-Bonferroni correction.

**Figure S3.**
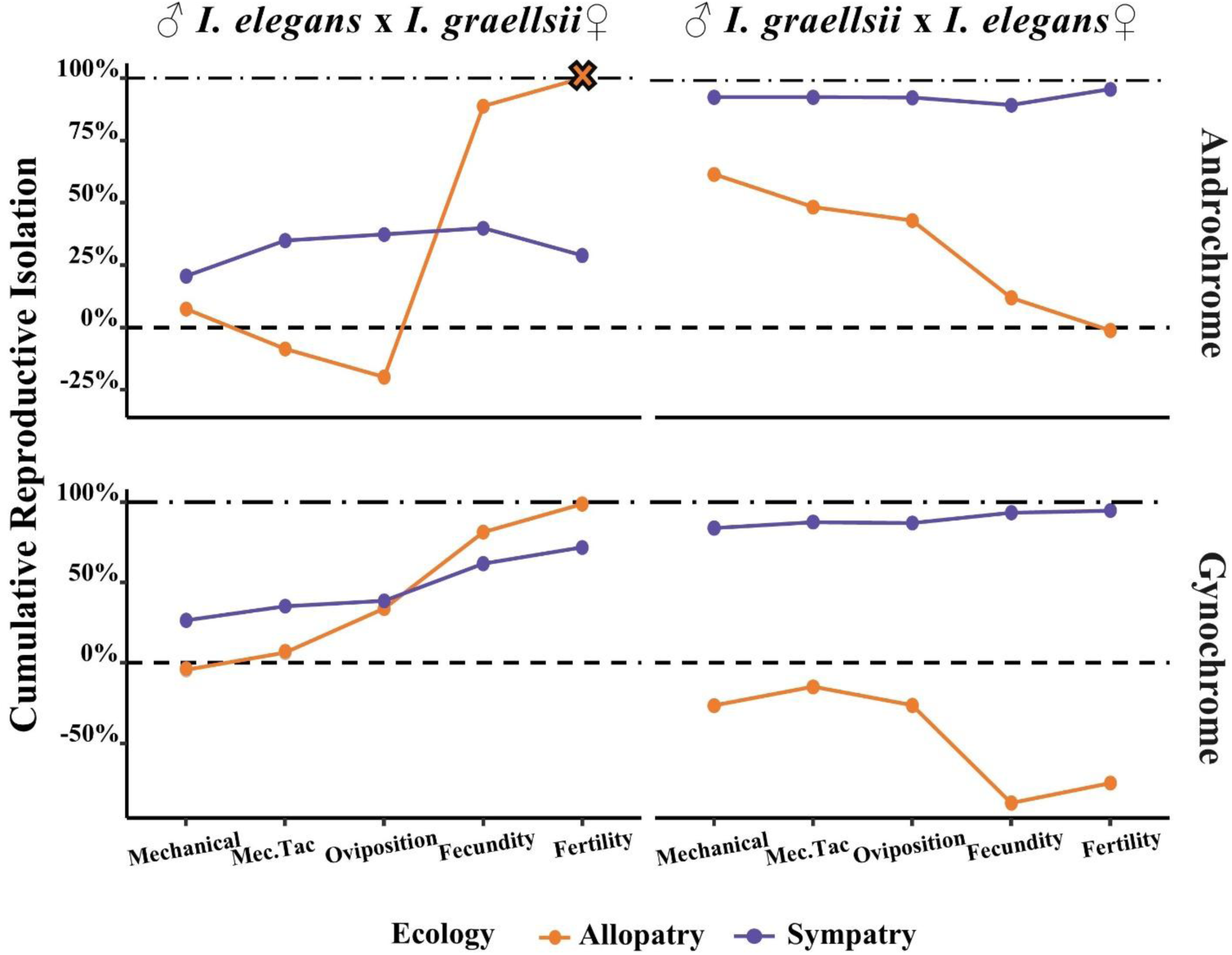
Cumulative isolation in both reciprocal crosses between *I. elegans* and *I. graellsii* in allopatry and sympatry. Upper panels display cumulative reproductive isolation for androchrome females, while lower panels show cumulative reproductive isolation for gynochrome females in allopatry (orange dots and line), and sympatry (purple dots and line). A cross indicates complete reproductive isolation and that the lack of gene flow between species in that cross direction. Horizontal dashed line indicates the threshold between reproductive isolation (above the line) and hybridization (below the line).

**Figure S4.**
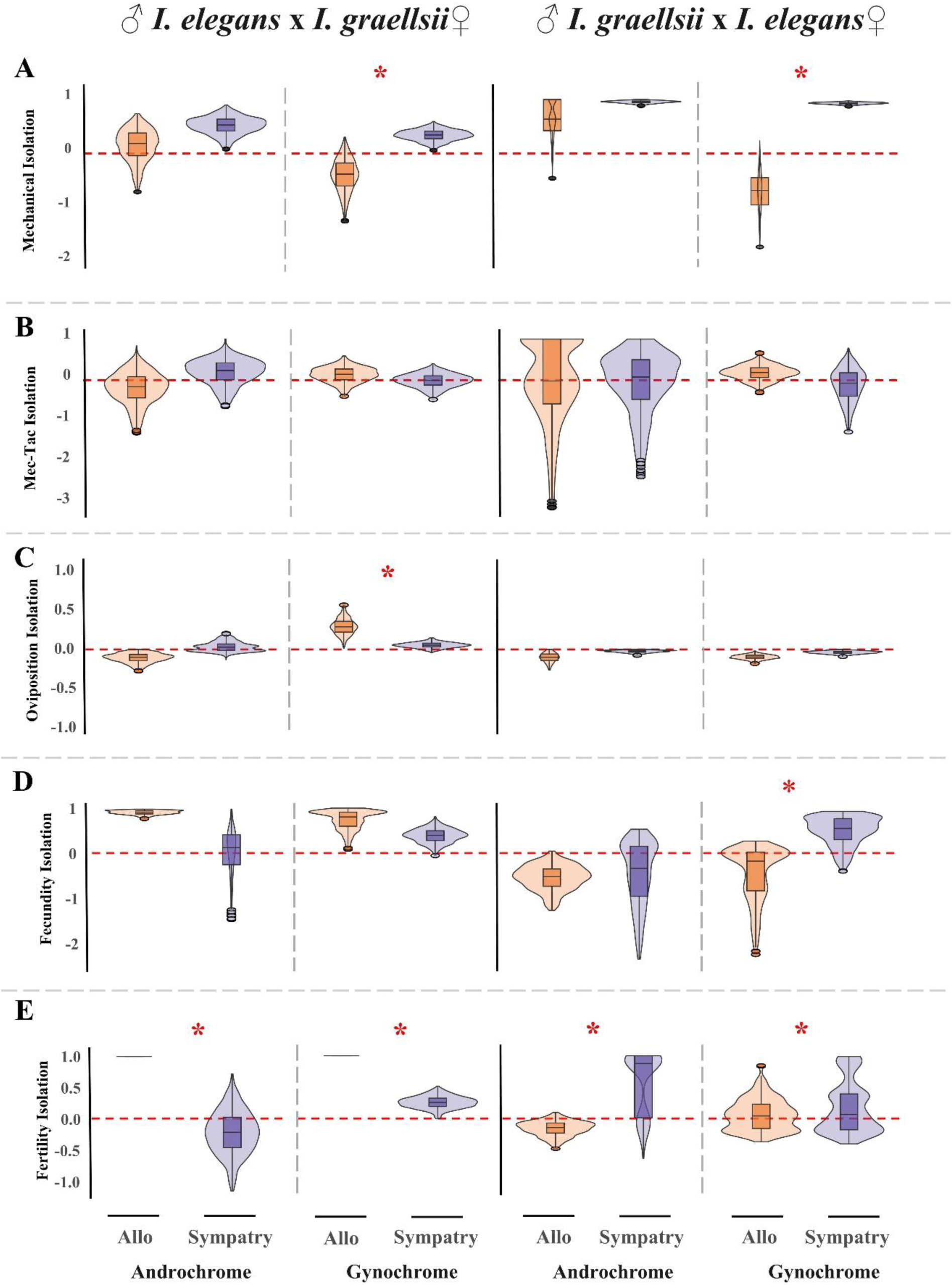
Absolute strength of each reproductive barrier and 95%CI between allopatry and sympatry. **A)** mechanical isolation, measured as the number of tandem attempts per heterospecific pair, **B)** mechanical-tactile isolation measured as the number of mating attempt per heterospecific pair, **C)** oviposition isolation, measured as the number of females that oviposited after mating. **D)** gametic isolation, fecundity, measured as the number of eggs laid by each female, and **E)** gametic isolation, fertility, measured as the ratio of fertile eggs laid by each female. Each barrier is presented with 95%CI estimated from 10,000 bootstrap resample of the dataset. The horizontal red dashed line marks the boundary between reproductive isolation (above the line) and facilitated hybridization (below the line). Red asterisks indicate statistically significant difference between the allopatric (orange boxplot and violin), and sympatric zone (boxplot and violin) after Holm-Bonferroni correction analyzed with the raw data.

**Figure S5.**
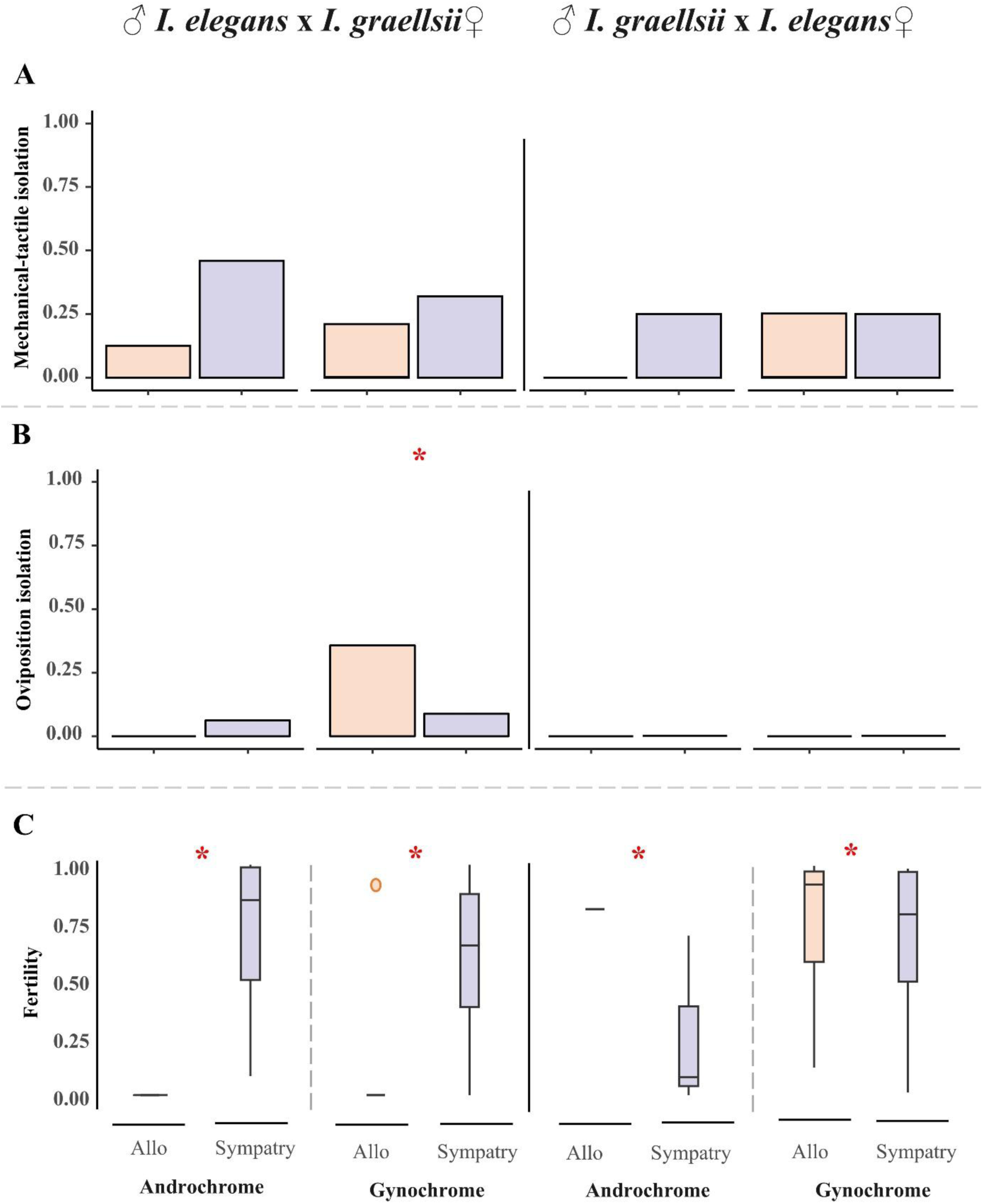
Differences in prezygotic barriers between allopatry and sympatry. **A)** Mechanical-tactile isolation, measured as the number of mating attempts per heterospecific pair. **B)** oviposition isolation, measured as the number of females that oviposited after mating. **C)** fertility, measured as the ratio of fertile eggs laid by each female. Red bold letters above the barplot, and boxplot indicate statistically significant difference the allopatric (orange barplot and boxplot), and sympatric zone (purple barplot and boxplot) after Holm-Bonferroni correction.

